# PIF7 controls leaf cell proliferation through an AN3 substitution-repression mechanism

**DOI:** 10.1101/2021.05.31.446395

**Authors:** Ejaz Hussain, Andrés Romanowski, Karen J. Halliday

## Abstract

Plants are agile, plastic organisms, able to adapt to ever-changing circumstances. Responding to far-red (FR) wavelengths from nearby vegetation, shade-intolerant species elicit the adaptive Shade Avoidance Syndrome (SAS), characterised by elongated petioles, leaf hyponasty and smaller leaves. We utilised end-of-day FR (EODFR) treatments to interrogate molecular processes that underlie the SAS leaf response. Genetic analysis establishes PHYTOCHROME INTERACTING FACTOR 7 (PIF7) is required for EODFR-mediated constraint of leaf blade cell division, while EODFR mRNAseq data identified *ANGUSTIFOLIA3 (AN3)* as a potential PIF7 target. We show PIF7 can suppress *AN3* transcription through a sequestering mechanism that prevents AN3 activation of its own expression. We also establish PIF7 and AN3 impose antagonistic control of gene expression via common cis-acting promoter motifs in several cell cycle regulator genes. EODFR triggers the molecular substitution of AN3 to PIF7 at G-box/PBE-box promoter regions, and a switch from promotion to repression of gene expression.

## Introduction

Growth plasticity is a fundamental property of plants, enabling adjustment to changes in the environment. The Shade Avoidance Syndrome (SAS) is a well-known adaptive response to the presence of nearby vegetation, that is characterised by gross changes in plant architecture and biomass (Sessa et al., 2018; Yang et al., 2016; Krahmer et al., 2021). A common feature of SAS is the reduction in leaf blade growth which can be dramatic in heavy vegetation shade. Our recent research, and that of others, has shown that phytochrome, an important modulator of the SAS leaf response, operates by controlling cell proliferation and expansion phases of development (Patel et al., 2013; Carabelli et al., 2018; Romanowski et al.; 2021). The molecular mechanisms through which phytochrome controls leaf growth are, however, unknown.

In Arabidopsis, phytochromes comprise a small gene family (PHYA-E) of photochromic biliproteins that are tuned to detect far-red (FR) light-rich conditions which occur in vegetation dense habitats (Franklin and Quail, 2010; Klose et al., 2020). The photo-isomeric properties of phytochrome dictate that red (R) light wavelengths drive the photoconversion from the inactive Pr to the active Pfr isomeric form. Exposure to FR wavelengths reverses this process, switching phytochrome to its inactive Pr state. Several studies have shown that FR inactivation of phyB-E, set in motion a series of molecular signalling events that activate SAS (Franklin and Quail. 2010). Although SAS is principally regulated by phyB, phyC has a contributory role, while phyD and phyE operate redundantly with phyB in this response (Franklin et al., 2004; Franklin and Quail. 2010). In contrast, phyA signalling, which is enhanced by periods of FR, acts to limit the extent of SAS, which can be detrimental if left unchecked (Yanovsky et al., 1995).

PhyB is known to operate by negatively regulating the PHYTOCHROME INTERACTING FACTOR (PIF) class of bHLH transcription factors. Studies indicate phyB can inhibit PIF1 and PIF3 action through sequestration, and can also initiate phosphorylation-mediated proteasomal degradation of PIF1, PIF3, PIF4 and PIF5 (Park et al., 2012, 2018; Pham et al., 2018). Consequently, deactivation of phyB by FR light leads to PIF de-repression and activation of transcriptional events. Of these PIFs, PIF4 and PIF5 have prominent roles in SAS, alongside another family member, PIF7, which has somewhat distinct regulatory characteristics. For instance, the phosphorylated form of PIF7 is not degraded, rather it is retained in the cytosol through interaction with 14-3-3 proteins (Huang et al., 2018). FR-rich shade light induces PIF7 de-phosphorylation and nuclear accumulation.

An important SAS feature, is the physiological response to FR is gated by the circadian clock, with a peak in FR responsiveness at dusk (Salter et al., 2003; Mizuno et al., 2015). This property means that end-of-day FR (EODFR) treatments can be relatively effective in eliciting SAS (Franklin, 2008; Romanowski et al., 2021). Further, delivery of short EODFR treatments avoids activating the SAS suppressor, phyA, assisting the delivery of a robust SAS response (Franklin and Quail 2010; Strasser et al., 2010). Recent reports have shown that PIF7 has a prominent role in mediating SAS responses induced by EODFR and that its action is clock gated (Salter et al., 2003; Mizuno et al., 2015; Leivar et al., 2020). FR shade light leads to the rapid activation of PIF7 through dephosphorylation and nuclear accumulation (Li et al., 2012; Huang et al. 2018; Leivar et al., 2020). These distinct molecular properties mean PIF7-mediated SAS signalling can be swiftly deployed following phyB deactivation at dusk, particularly in short days when PIF7 is reported to be most effective (Leivar et al., 2020). After an EODFR treatment, PIF7 is primarily active post-dusk, as rising levels of the night-phased clock component ELF3 gradually suppress PIF7 action, through direct interaction and prevention of PIF7 DNA-binding (Jiang et al., 2019).

Earlier studies demonstrated that simulated canopy shade can constrain the phase of leaf cell proliferation in a process involving HD-Zip II transcription factors *ATHB2* and *ATHB4* (Carabelli et al., 2007; Carabelli et al., 2018). Our recent mRNAseq analysis identified key leaf development genes as EODFR regulated, opening a potentially novel phytochrome signalling route (Romanowski et al., 2021). Amongst these genes, *ANGUSTIFOLIA 3/GRF-INTERACTING FACTOR 1 (AN3/GIF1) GROWTH REGULATING FACTORs GRF2, GRF4* and *GFR6*, and *BRAHMA (BRM)* are known to regulate leaf blade cell proliferation, while, the small, narrow leaf phenotype of the *an3-4* mutant is reminiscent of the *phyB* leaf blade phenotype (Tsukaya et al., 2002; Kim and Kende, 2004, Horiguchi et al., 2005, Vercruyssen et al., 2014). AN3 is proposed to operate centrally in a complex with DNA-binding GRFs and with SWITCH/SUCROSE NONFERMENTING (SWI/SNF) chromatin remodelling proteins, like BRM and BAF60, to regulate transcription (Vercruyssen et al., 2014). Interestingly the AN3-GRF-SWI/SNF complex is highly conserved in eudicots and monocots and therefore represents a widespread leaf development mechanism (Nelissen et al., 2015; Besbrugge et al., 2018; Shimano et al., 2018).

This study elucidates a novel molecular mechanism through which EODFR suppresses leaf growth. We demonstrate that PIF7 is a potent inhibitor of leaf cell proliferation following SAS-inducing EODFR treatments. Genetic analysis indicates this is accomplished through the repression of AN3. We show PIF7 suppresses *AN3* expression through a sequestration mechanism that prevents AN3 self-activation. Our data also point to PIF7 and AN3 signalling convergence at common promoter cis-elements in cell cycle genes. EODFR induces PIF7 substitution for AN3 at target promoters and a concomitant shift from promotion to the repression of gene expression.

## Results

### EODFR represses cell division during two phases during leaf blade development

Previously we used phytochrome-deactivating EODFR treatments to mimic and test the impact of vegetative shading at different phases of rosette leaf 3 (L3) development (Romanowski et al., 2021). We established that daily EODFR was effective in limiting L3 expansion by suppressing epidermal cell division or expansion, contingent on whether EODFR coincided with the proliferation or expansion phase of development. Extending these findings we found that EODFR treatment delivered daily (light: dark (LD) 12-h: 12-h photoperiod at 22 °C), from day 6, or *phyB-9*, are equally effective in reducing the cell number, but not the size of both epidermal and palisade cells (Supplementary Fig. 1a-i). We observed similar EODFR effects on leaf blade expansion for all rosette leaves (Supplementary Fig. 2) but focus on L3 as a model to allow direct comparison with our published data and that of other studies (Kozuka et al., 2005; Andriankaja et al., 2012; Beltramino et al., 2018; Romanowski et al., 2021).

To pinpoint more precisely the time-window during which phytochrome controls blade cell division we exposed plants grown in light: dark (LD) 12-h: 12-h photoperiod at 22 °C, to 5 days of EODFR in the following intervals: day 6-10, 10-14 or 14-18, negative controls did not receive EODFR, and positive controls received EODFR for day 6-18, or 6-34 (Fig.1a). As expected, EODFR delivered for the longest period, 6-34 days, was the most effective in reducing blade area and limiting cell division, followed by the 6-18 interval (Fig.1b-h). EODFR provided in the 6-10 or 14-18 time-slots did not induce significant changes in leaf size or cell division, in contrast to the 10-14 day window, which was as effective as the longer 6-18 period. The data identify two windows during which phyB controls cell division: day 10-14, which coincides with the main proliferative phase, and day 6-34 which probably also includes the meristemoid cell division phase (Gonazalez et al., 2012).

**Fig. 1:**
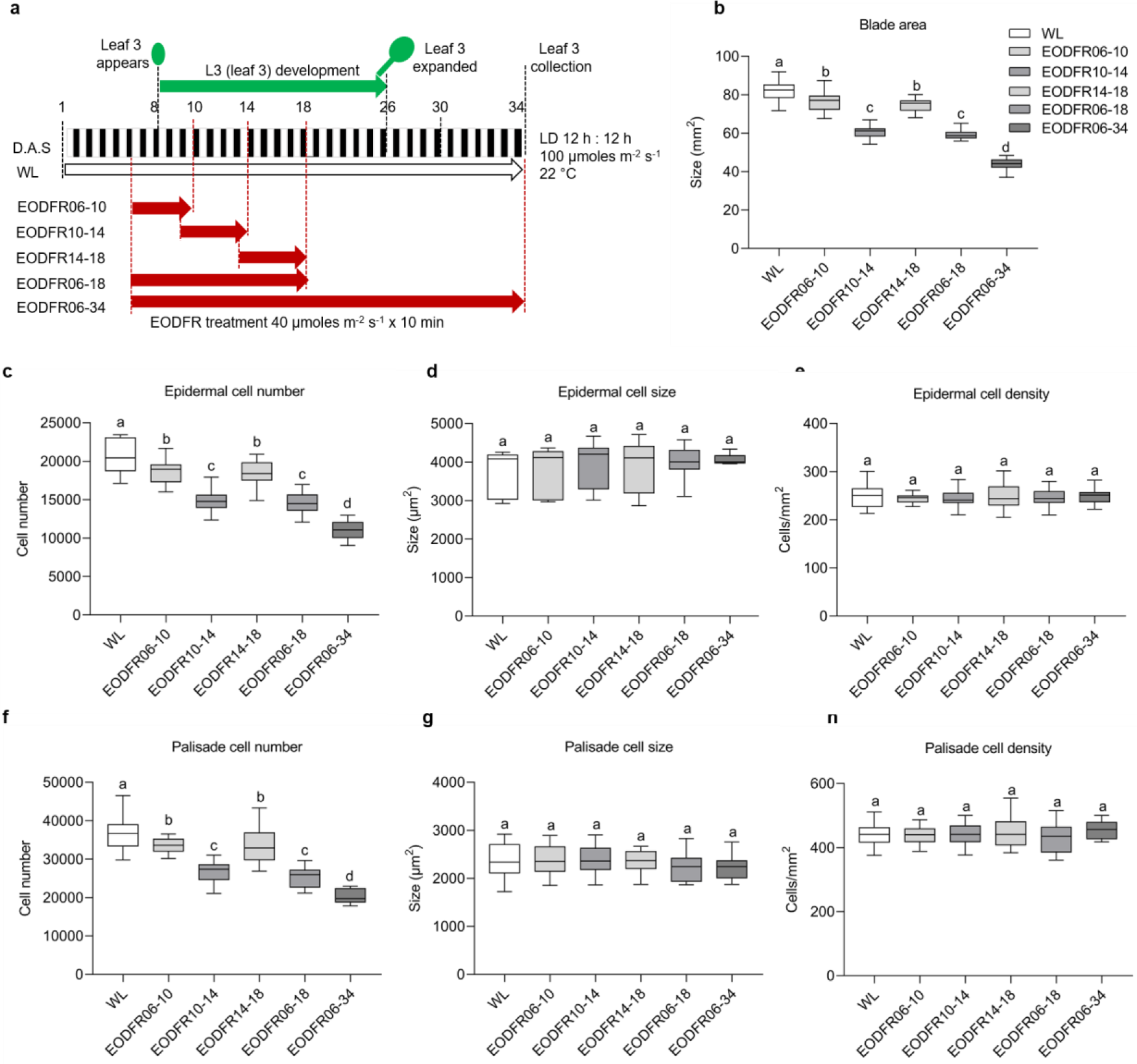
EODFR controls leaf expansion by restricting two phases of cell division. **a** Schematic representation of the environmental treatment regime. White and black rectangles indicate the 12 h of day or night period, respectively. The green arrow indicates the period of L3 (leaf 3) development, which is enclosed between the two innermost black dashed lines. The plant drawn on top of day 8 indicates L3 emergence. The leaf drawn on top of day 26 indicates that L3 is fully expanded. The red dashed lines indicate the day at which a specific treatment was started and coincides with a specific coloured arrow (WL in white and EODFR in dark red). The red dashed line at the end of day 10, 14, 18 and 34 marks the end of each treatment. The black dashed line on day 34 indicates leaf 3 collection for each treatment. **b** Comparison of leaf blade area (ANOVA, Tukey’s post hoc test, *p <* 0.0001, *n >* 20 blades). **c**, **f** epidermal and palisade cell number (ANOVA, Tukey’s post hoc test, *p <* 0.0001 in (**c**, **f**), *n >* 20 blades), **d**, **g** epidermal and palisade cell size (ANOVA, Tukey’s post hoc test, *p* = 0.4234 in (**d**) and *p* = 0.375327 in (**g**); 27 cells per blade, *n >* 20 blades), **e**, **h** epidermal and palisade cell density (ANOVA, Tukey’s post hoc test, *p* = 0.8082 in (**e**) and *p* = 0.7436 in (**h**), *n >* 20 blades). **b**, **c**, **d**, **e**, **f**, **g**, **h** Error bars represent the SEM. The center of the error bars represents the mean values. Different letters denote statistically significant differences in leaf blade area, cell number, cell size and cell density between different treatments (ANOVA followed by Tukey’s post hoc test). This experiment was repeated at least two times with similar results.

### mRNAseq identifies EODFR regulated genes implicated in cell cycle control

As the molecular connections between phyB and cell cycle control are largely unknown wesought to identify candidate cell cycle regulator genes from our L3 mRNAseq data (Romanowski et al., 2021). This longitudinal study captured EODFR-induced changes in transcription through leaf development. From this dataset, we identified 317 genes, classified as transcription factors, or co-transcriptional regulators (Supplementary Table 1), with EODFR-altered expression early-on in L3 development. In this gene set, 14 genes (*AN3, GRF2, GRF4, GRF6, DP-E2FA like1 (DEL1), AINTEGUMENTA, (ANT), DP-E2F like3 (DEL3), FAMA, (FMA), SCARECROW (SCR), AT4G02110, HOMEOBOX GENE 8 (HB-8), AUXIN RESPONSE FACTOR 10 (ARF10), ASYMMETRIC LEAVES 1 (AS1)* and *MYB DOMAIN PROTEIN 3R-4 (MYB3R)* are known to have roles associated with the cell cycle (Kim and Kende, 2004; Lee et al., 2009; Van Leene et al., 2010). In nearly all cases, expression levels were higher on day 13, which falls in the 10-14 day window that coincides with cell proliferation in our experimental regime (Supplementary Fig. 3a-c) (Romanowski et al., 2021; https://aromanowski.shinyapps.io/leafdev-app/).

Recent articles have identified AN3 as a transcription regulator with a central role in the promotion of leaf expansion (Horiguchi et al., 2005; Kawade et al., 2013; Liebsch et al., 2020). AN3 lacks DNA-binding capacity, and so functions as a transcriptional co-activator by interacting with GRFs that can bind DNA directly (Kim et al., 2012; Vercruyssen et al., 2014; Nelissen et al., 2015; Besbrugge et al., 2018; Omidbakhshfard et al., 2018). The AN3-GRF unit appears to regulate transcription by recruiting chromatin modelling components, including *BRM,* which operates as a central SWI2/SNF2 ATPase (Vercruyssen et al., 2014). Our finding opens the possibility that the AN3 transcriptional module operates downstream of phytochrome to control leaf cell division and growth.

### EODFR regulation of cell division is abolished in the *an3-4* mutant

To establish whether AN3 is implicated in phytochrome control of the cell cycle, we first analysed the EODFR leaf response in the *an3-4* mutant. In line with earlier reports (Horiguchi et al., 2005; Kawade et al., 2013), in our conditions, *an3-4* L3 blades were smaller than wild type (WT) with fewer cells (Fig. 2a, b and Supplementary Fig. 4b). We found EODFR was less effective in reducing the blade area of *an3-4* compared to WT, but *an3-4* was still EODFR responsive (Fig. 2a). In contrast, while EODFR markedly reduced WT epidermal and palisade cell number, *an3-4* was completely insensitive to the treatment (Fig. 2b and Supplementary Fig. 4b). We also established that, similar to earlier studies (Horiguchi et al., 2005; Kawade et al., 2013), in our control conditions *an3-4* epidermal and palisade cell size were slightly larger than WT (Supplementary Fig. 4a, c, d, e). However, EODFR application completely suppresses the cell size defect caused by the *an3-4* mutation (Supplementary Fig. 4. a, c) This finding most likely accounts for the retention of an *an3-4* blade area EODFR response. In summary, our data show *an3-4* has a constitutively low cell number, which is insensitive to EODFR, which implies AN3 may operate downstream of phytochrome to promote leaf blade cell division. In addition, our data indicate that phytochrome inactivation by EODFR prevents cell expansion caused by the *an3-4* mutation.

**Fig. 2:**
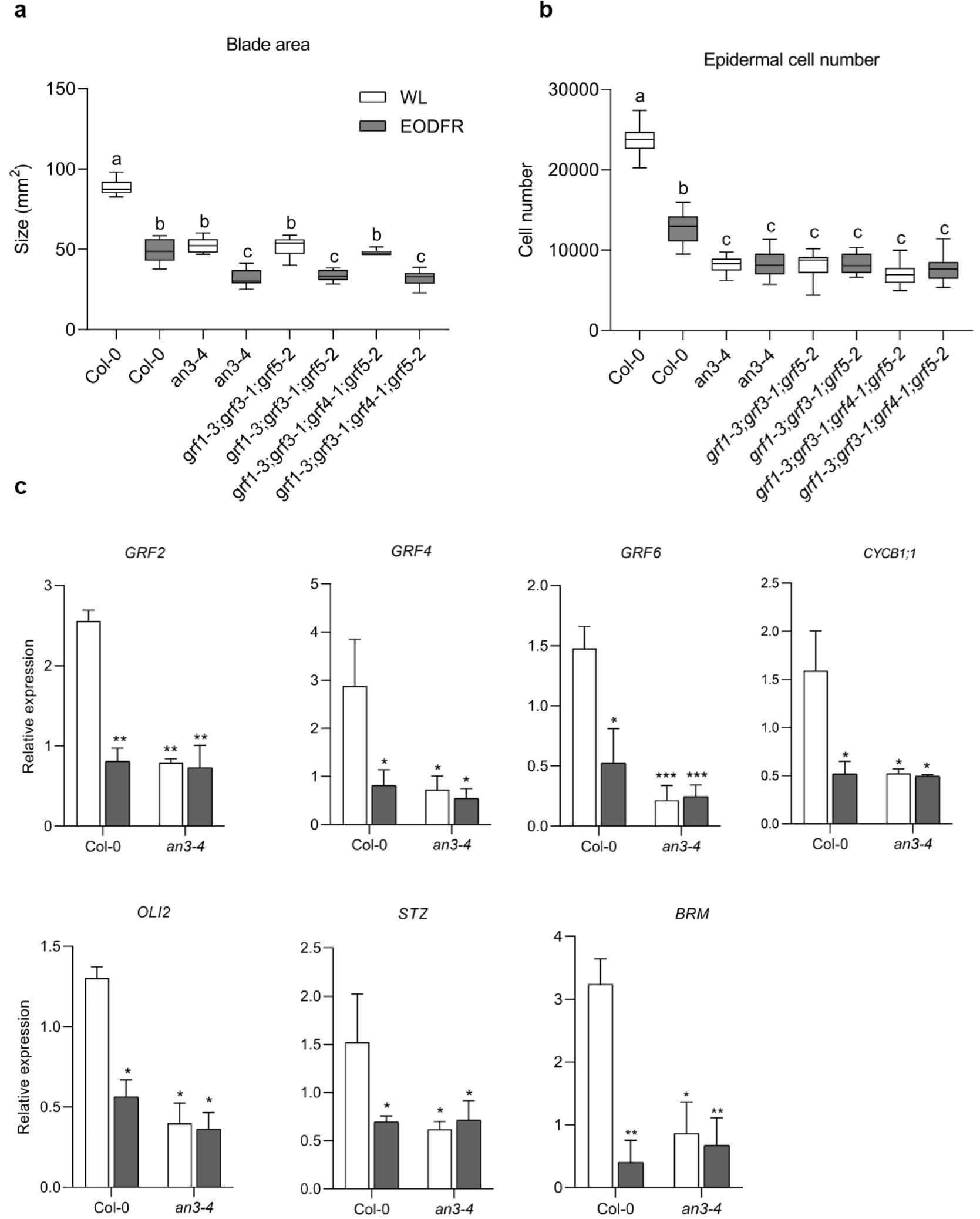
The *an3-4* mutant is completely insensitive to EODFR regulation of leaf cell division. **a** Comparison of leaf blade area (ANOVA, Tukey’s post hoc test, *p <* 0.0001, *n =* 15 blades). **b** Epidermal cell number (ANOVA, Tukey’s post hoc test, *p <* 0.0001, *n =* 15 blades). **a**, **b** Error bars represent the SEM. The center of the error bars represents the mean values. Different letters denote statistically significant differences in leaf blade area and cell number between genotypes and different treatments (ANOVA followed by Tukey’s post hoc test). This experiment was repeated at least two times with similar results. **c** *GRF2, GRF4, GRF6, CYCB1;1, OLI2, STZ* and *BRM* messenger RNA (mRNA) level using quantitative reverse transcription (RT-qPCR) after the shift from white light to EODFR in Col-0 and *an3-4*. Seedlings were grown for 13 days under a light : dark (LD) 12 h : 12 h photoperiod, at 22 °C. On day 13, seedlings were either shifted to EODFR treatment or kept in the white light control condition. L3 blades were harvested 13 days after sowing at zeitgeber (ZT) 24. The transcript levels were calculated relative to those of *PP2A*. Error bars represent the s.d. of three biological replicates. The center of the error bars represents the mean values. (\**p* < 0.05, \*\**p* < 0.01 and \*\*\**p* < 0.001, Student’s *t*-test).

Since AN3 complexes with GRFs, which are reported to have redundant roles in regulating leaf size, we analysed the EODFR responsiveness of multi-allele *grf* mutants (Kim and Kende, 2004). We found that the triple *grf1-3;grf3-1;grf5-2* and quadruple *grf1-3;grf3-1;grf4-1;grf5-2* mutants are phenotypically similar to *an3-4*, with small leaf blades, low epidermal and palisade cell numbers, and reduced sensitivity to EODFR (Fig. 2a,b and Supplementary Fig. 4a-e). These data, therefore, strengthen the hypothesis, that phytochrome control of leaf cell proliferation is mediated, at least in part, through the AN3-GRF transcriptional complex.

### AN3 is required for phyB-controlled expression of leaf cell cycle regulators

Several of the leaf developmental genes identified as EODFR repressed in our mRNAseq dataset are known AN3 targets (Lee et al., 2009; Vercruyssen et al., 2014; Besbrugge et al., 2018; Romanowski et al., 2021). For instance, *Chromatin Immunoprecipitation (ChIP) qPCR* has shown AN3 can directly associate with the *GRF6* promoter, while *GRF2*, *GRF4*, *OLIGOCELLULA 2* (*OLI2*), *SALT TOLERANCE ZINC FINGER* (STZ), and *BRM* genes were shown to be bound by AN3 in tandem chromatin affinity purification sequencing (TChAP-seq) and chromatin affinity purification sequencing (ChIP-seq) (Vercruyssen et al., 2014; Besbrugge et al., 2018). Further, overexpression of *AN3* resulted in increased expression of *CYCB1;1,* identifying this cell cycle gene as AN3 regulated (Lee et al., 2009; Vercruyssen et al., 2014). We reasoned that if the phytochrome-controlled cellular response is mediated through AN3 then EODFR control of these AN3 targets would be curtailed in the *an3-4* mutant. Indeed, compared to WT, our quantitative reverse transcriptase PCR (qRT-PCR) assay showed that this gene-set has reduced expression in *an3-4* with a diminished or lack of response to EODFR (Fig. 2c). This was also the case for the AN3 targets *GRF1, GRF3* and *GRF5,* and key cell cycle genes *CDC45*, *CDC6,* and *DEL1,* identified as EODFR-repressed in our mRNAseq data (Supplementary Fig. 3d) (Romanowski et al., 2021), but not known to be AN3 regulated (Supplementary Fig. 4f) (Vercruyssen et al., 2014). These data identify *CDC45*, *CDC6,* and *DEL1* as AN3 regulated, and support the hypothesis that AN3 action is dependent on phytochrome. Our results suggest AN3 promotes the expression of genes that modulate the leaf cell cycle in control conditions, but not in EODFR which deactivates phyB and other SAS controlling phytochromes.

### AN3 operates downstream of PIF7 to control leaf cell division

PIF7 is known to have an important function in EODFR-induced seedling and adult plant responses, while the contribution from other PIFs is smaller (Mizuno et al., 2015; Xie et al., 2017; Leivar et al., 2020). Consistent with this notion, we show *pifq,* which lacks PIF1, PIF3, PIF4 and PIF5, has a smaller leaf blade area with fewer epidermal cells in control conditions but still responds to EODFR, although the response is reduced (Supplementary Fig. 5a-d). In contrast, *pif7-1* has a WT leaf blade size and is completely unresponsive to EODFR (Fig. 3a). Likewise, the *pif7-1* mutant has a similar epidermal cell number and cell size to WT white light control and lacks the EODFR reduction in cell number (Fig. 3b-d). These data reflect the known functional properties of PIF7 which is activated by EODFR and additionally illustrate that PIF7 is necessary for the EODFR suppression of phytochrome‐mediated leaf cell proliferation (Leivar et al., 2020).

**Fig. 3:**
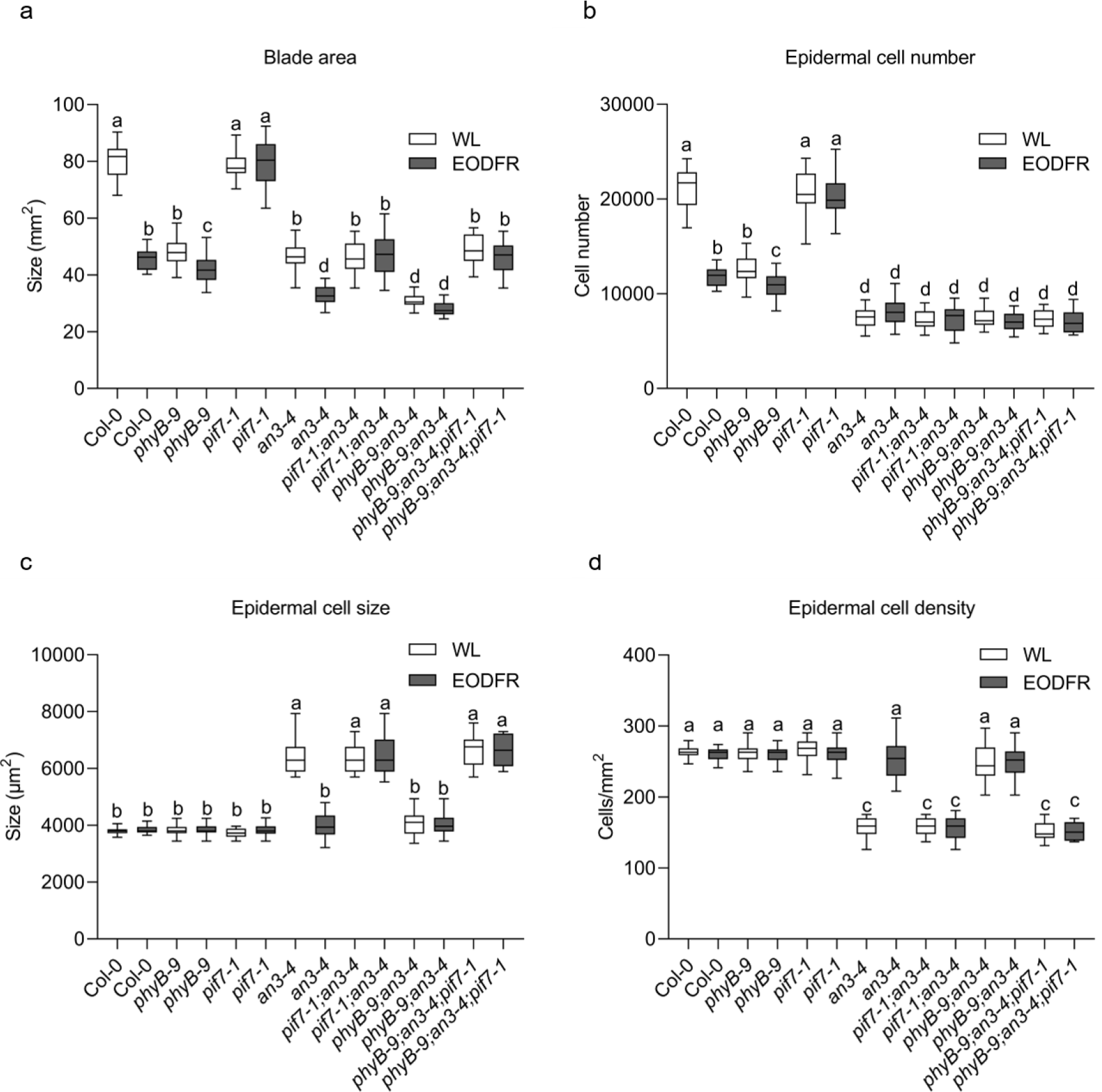
Both *phyB-9* and *pif7-1* are epistatic to *an3-4*. **a** Comparison of leaf blade area (ANOVA, Tukey’s post hoc test, *p <* 0.0001, n > 20 blades). **b** epidermal cell number (ANOVA, Tukey’s post hoc test, *p <* 0.0001, *n >* 20 blades), **c** epidermal cell size (ANOVA, Tukey’s post hoc test, *p <* 0.0001 in (**d**); 27 cells per blade, n > 20 blades), **d** epidermal cell density (ANOVA, Tukey’s post hoc test, *p <* 0.0001, n > 20 blades). Plants were grown under a light : dark (LD) 12 h : 12 h photoperiod, at 22 °C. On the 6th day, the plants were exposed to EODFR treatment, until the 34th day. All leaf 3 blades were harvested and measured on the 34th day. **a**, **b**, **c**, **d** Error bars represent the SEM. The center of the error bars represents the mean values. Different letters denote statistically significant differences in leaf blade area, cell number, cell size and cell density between genotypes and different treatments (ANOVA followed by Tukey’s post hoc test). This experiment was repeated at least two times with similar results.

As we have shown that AN3 promotion of cell proliferation is prevented by EODFR, our data implicate PIF7 as a negative regulator of AN3. To begin to test this hypothesis we generated the *an3-4;phyB-9*, *an3-4;pif7-1* and *an3-4;phyB-9*;*pif7-1* multi-allele mutants. Fig.3a,b shows, that, as expected, *phyB-9* L3 blades are smaller and narrower than those of WT with reduced epidermal cell number in control conditions, while *pif7-1* lacks WT response to EODFR. Comparison of cell number in *an3-4;phyB-9*, *an3-4;pif7-1, an3-4;phyB-9*;*pif7-1* and the single mutant parental lines revealed *an3-4* epistasis over *phyB-9,* which was most evident in control conditions, and *an3-4* epistasis over *pif7-1,* particularly under EODFR. Thus, AN3 appears to be required for phyB-PIF7 module control of leaf cell division.

With regards to L3 epidermal cell size, *phyB-9* and *pif7-1* are indistinguishable from the WT, and as shown earlier, *an3-4* cells are larger in control but not in EODFR conditions (Fig.3c-d). In contrast to cell number, for cell size *phyB-9* is epistatic to *an3-4* in control conditions, and *pif7-1* is epistatic to *an3-4* in EODFR, indicating, *an3-4* cell expansion is dependent on whether phyB is active. Further, *an3-4;phyB-9*;*pif7-1* is indistinguishable from *an3-4:pif7-1* (Fig.3a), which is consistent with PIF7 operating downstream of phyB. Collectively, our data indicate that AN3 is required for phyB promotion of cell division in control conditions. EODFR induced PIF7 activation, restricts cell division, possibly by repressing AN3 levels and/or activity, and secondarily, PIF7 appears to limit the compensatory cell expansion that arises from AN3 deactivation.

### PIF7 suppresses *AN3* expression through direct binding to the *AN3* promoter

Our mRNAseq data showed *AN3* transcript abundance is lowered by EODFR, and our genetic data identified PIF7 as a potential *AN3* regulator (Romanowski et al., 2021). To test this, we quantified *AN3* transcript levels in *pif7-1* by qRT-PCR assay. Figure 4a shows *AN3* transcript levels are unaltered by EODFR in *pif7-1*, which implicates PIF7 as a repressor of *AN3* transcription. A recent study reported that as for other PIFs, PIF7 preferentially binds to G-boxes (CACGTG) and PBE-boxes (CA[TG/CA]TG) in the promoters of target genes (Galvao et al., 2019). Interestingly, in Arabidopsis, a G-box motif has been identified as an important transcriptional cis-regulatory element in the *AN3* promoter, thus we surmised PIF7 may directly regulate *AN3* transcription (Liu et al., 2020). To test this we, performed ChIP-qPCR assays using 13-day-old leaf tissues of WT and PIF7-Flash (*35S::PIF7-Flash*; 9xMyc-6xHis-3xFlag) plants (Fig. 4c; Li et al., 2012). This assay showed PIF7-Flash enrichment at the G-box containing region of the *AN3* promoter (P1), which was further enhanced after EODFR (Fig. 4b). We also recorded PIF7-Flash enrichment at the G-box containing promoter region of the known PIF7 target, *IAA19*, but not in either of the negative controls, a fragment of AN3 CDS region (P4) and no-antibody ChIP tissues (Fig. 4b,d) (Peng et al., 2018). These observations together with the *AN3* expression data, (Fig. 4a) indicate that PIF7 directly represses *AN3* expression through association with the G-box region (P1) of the *AN3* promoter.

**Fig. 4:**
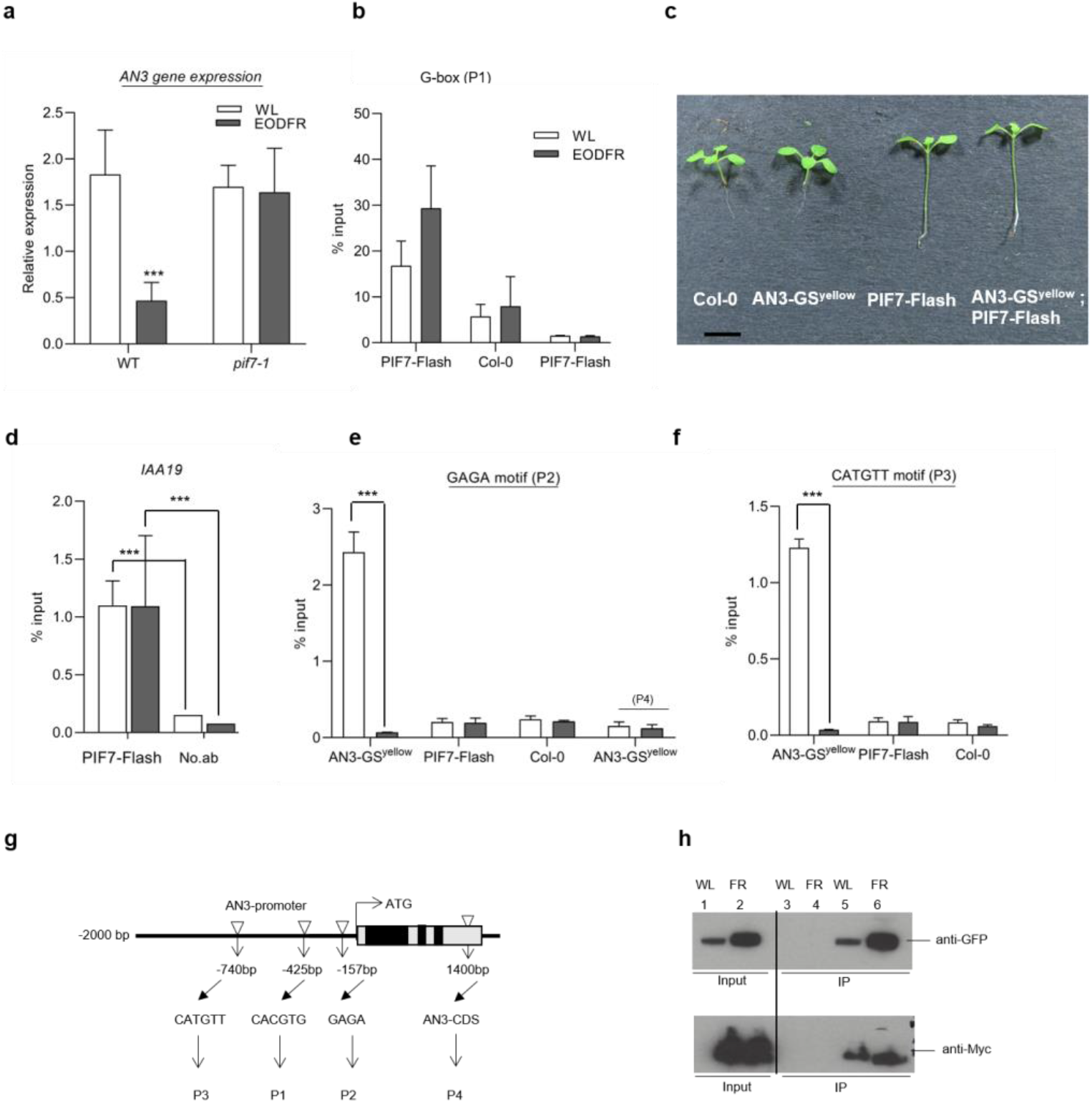
PIF7 directly suppress AN3 expression to restrict leaf cell division. **a** *AN3* messenger RNA (mRNA) level using quantitative reverse transcription (RT-qPCR) in Col-0 and *pif7-1* seedlings grown for 13 days under a light : dark (LD) 12 h : 12 h photoperiod, at 22 °C. On day 13, seedlings were either shifted to EODFR treatment or kept in the white light control condition. L3 blades were harvested 13 days after sowing at zeitgeber (ZT) 24. The transcript levels were calculated relative to those of *PP2A*. Error bars represent the s.d. of three biological replicates. The center of the error bars represent the mean values. (\**p* < 0.05, \*\**p* < 0.01 and \*\*\**p* < 0.001, Student’s *t*-test). **b, d, e, f** ChIP assay showing enrichment of 35S::PIF7-Flash and 35S::AN3-GS^yellow^ on *AN3* fragments (P1-P4) and *IAA19* promoter. Seedlings were grown as described in (**a**) samples were taken at ZT14. The wild type (Col-0) plants acted as a control. Error bars represents the s.d. of three biological replicates. The center of the error bars represent the mean values. (\**p* < 0.05, \*\**p* < 0.01 and \*\*\**p* < 0.001, Student’s *t*-test). **c**Representative 13-day-old Col-0, 35S::AN3-GSyellow, 35S::PIF7-Flash and 35S::PIF7-Flash;35S::AN3-GS^yellow^ seedlings grown as described in (**a**). Bar = 5 mm^2^. **g** The schematic diagram of AN3 promoter and gene body showing P1-4 fragments. **h** co-immunoprecipitation experiment showing the interaction between AN3 and PIF7 in vivo. Seedlings were grown as described in (**a**), samples were taken at ZT14. The 35S::AN3-GS^yellow^ and 35S::PIF7-Flash;35S::AN3-GS^yellow^ were precipitated with an anti-Myc antibody. 35S::AN3-GS^yellow^ and 35S::PIF7-Flash input and precipitated fractions were detected by immunoblots using anti-GFP and anti-Myc antibodies, respectively. **1** and **2** Immunoblots showing the co-immunoprecipitation results from Input of 35S::PIF7-Flash;35S::AN3-GS^yellow^ from white light and EODFR samples, respectively. **3** and **4** Immunoblots showing the co-immunoprecipitation results from IP (immunoprecipitation) of 35S::AN3-GS^yellow^ from white light and EODFR samples, respectively. **5** and **6** Immunoblots showing the co-immunoprecipitation results from IP of 35S::PIF7-Flash;35S::AN3-GS^yellow^ from white light and EODFR samples, respectively.

### EODFR prevents AN3 binding to its own promoter

We reasoned that if PIF7 was operating through *AN3* to regulate leaf blade cell proliferation, then PIF7 would also be required for EODFR suppression of AN3 target genes. This is indeed what we observed, as for each of the AN3 regulated genes (*GRF1, GRF2, GRF3, GRF4, GRF5, GRF6, CYCB1;1, OLI2, STZ, BRM, CDC45, CDC6,* and *DEL1*) *pif7-1* completely abolished the EODFR repression response (Supplementary Fig. 6). Next, we tested whether PIF7 functions solely by regulating *AN3* expression by analysing a 35S::AN3-GS^yellow^ line (subsequently referred to as AN3-GS^yellow^, (Besbrugge et al., 2018). Consistent with published data we found that the AN3-GS^yellow^ line had elevated *AN3* expression and a moderately increased leaf blade area in control conditions (Supplementary Fig. 7a,b) (Besbrugge et al., 2018). However, this line exhibited a WT response to EODFR, suggesting PIF7 may not regulate AN3 solely by modulating its transcription, rather it might also regulate AN3 activity.

Published studies indicate AN3 can bind to its own promoter to promote expression through self-regulatory motifs (the GAGA-motif present in the region encoding the 5’ untranslated region (UTR) and a CATGTT box) in the *AN3* promoter (Vercruyssen et al., 2014; Meng et al., 2018) (Fig.4g). As PIF7 has not been shown to bind to these motifs, we used ChIP-qPCR to test whether PIF7 activation by EODFR disrupts AN3 association with self-regulatory motifs. As expected, we observed statistically significant enrichment of AN3-GS^yellow^ (p-values < 0.001) at the GAGA (P2) and CATGTT containing regions (P3), but not the AN3 coding sequence control (P4) (Fig.4e,f). Notably, we did not observe AN3-GS^yellow^ association with P2 or P3 following EODFR. We next tested if PIF7 could bind to these regions using PIF7-Flash expressing plants and found that, while we detected PIF7 enrichment at the promoter of a known target, *IAA19* (p-value < 0.001), PIF7 did not associate with regions containing the GAGA or CATGTT motifs in control or EODFR conditions (Fig. 4d-f). Thus, our data indicate that EODFR prevents AN3 association with self-regulatory elements, possibly through a PIF7-dependent mechanism.

To test this more rigorously we performed co-immunoprecipitation in lines co-expressing PIF7-Flash and AN3-GS^yellow^ (Fig. 4b). We found that AN3-GS^yellow^ co-immunoprecipitated with PIF7-Flash in both control and EODFR conditions (Fig. 4h). The interaction in control conditions is consistent with the increased levels and activity of PIF7, evidenced by the small L3 blade of the 35S::PIF7-Flash line, which was comparable to EODFR treated WT plants (Supplementary Fig. 7b). Interestingly, we also observed higher levels of AN3 protein in EODFR, which implies that either AN3 protein synthesis or degradation is altered by EODFR (Fig. 4h). Taken together, our results suggest EODFR conditions promote PIF7 action and sequestration of the AN3 complex from target leaf cell cycle gene promoters. This proposition is also consistent with the leaf blade phenotype of double overexpressing 35S::PIF7-Flash;35S::AN3-GS^yellow^ plants, in which PIF7 exhibits dominant action over AN3 (Supplementary Fig. 7b).

### EODFR induces a switch from AN3 to PIF7 at common target promoters

Close inspection of previously published AN3-HBH TChAPseq data revealed a high-confidence AN3 protein binding peak in gene promoter regions containing GAGA, PBE-box and G-box motifs and ChIP-qPCR assay confirmed AN3 binding on PBE-boxes of *GRF5* and *GRF6* promoters (Vercruyssen et al., 2014). Indeed, *cis-*acting regulatory DNA element analysis showed that all the AN3 regulated genes in this study possessed G-box (CACGTG) and/or PBE-box (CA[TG/CA]TG) promoter motifs (Supplementary Figure. 9). This finding provided the possibility that AN3 and PIF7 signalling may converge at these *cis-*elements in a common set of genes, and that PIF7 may interfere with AN3 association and/or activity at the target promoters. To explore this notion, we performed ChIP-qPCR assays using the AN3-GS^yellow^ and PIF7-Flash lines. As shown in Fig. 5a and Supplementary Fig. 8a, AN3-GS^yellow^ showed enrichment at G-box and or PBE-box promoter regions for *GRF2, GRF4, GRF6, CYCB1;1, OLI2, STZ, GRF1, GRF3, GRF5, CDC45, CDC6,* and *DEL1,* in control but not in EODFR conditions. In contrast, we recorded higher PIF7 enrichment in EODFR (p-values < 0.0001) at the same G-box or PBE-box containing promoter regions (Fig. 5b and Supplementary Fig. 8b). Again, control behaviour was as expected, with PIF7-Flash enrichment at the G-box promoter region of *IAA19*, but not the no antibody control. (Fig. 5e) (Peng et al., 2018). Similarly, we also observed AN3-GS^yellow^ enrichment at the GAGA-motif positive control and not at the AN3 CDS region where AN3 is not expected to associate (Fig. 5d). These data show that PIF7 and AN3 can associate with the G-box and PBE-box regions of common genes and that PIF7 promoter occupancy in EODFR may prevent AN3-binding. Our data are further corroborated by the *pif7-1;*AN3-GS^yellow^ ChIP-qPCR data which shows, in the absence of functional PIF7, AN3 remains enriched at G-box or PBE-box promoter regions (p-values < 0.0001) under EODFR as well as control conditions (Fig. 5c and Supplementary Fig. 8c). Collectively our analysis indicates that AN3 and PIF7 antagonistically regulate a common set of genes that control leaf cell proliferation. EODFR leads to the activation of PIF7 which dislodges and substitutes for AN3 at G-box or PBE-box containing regions of target promoters.

**Fig. 5:**
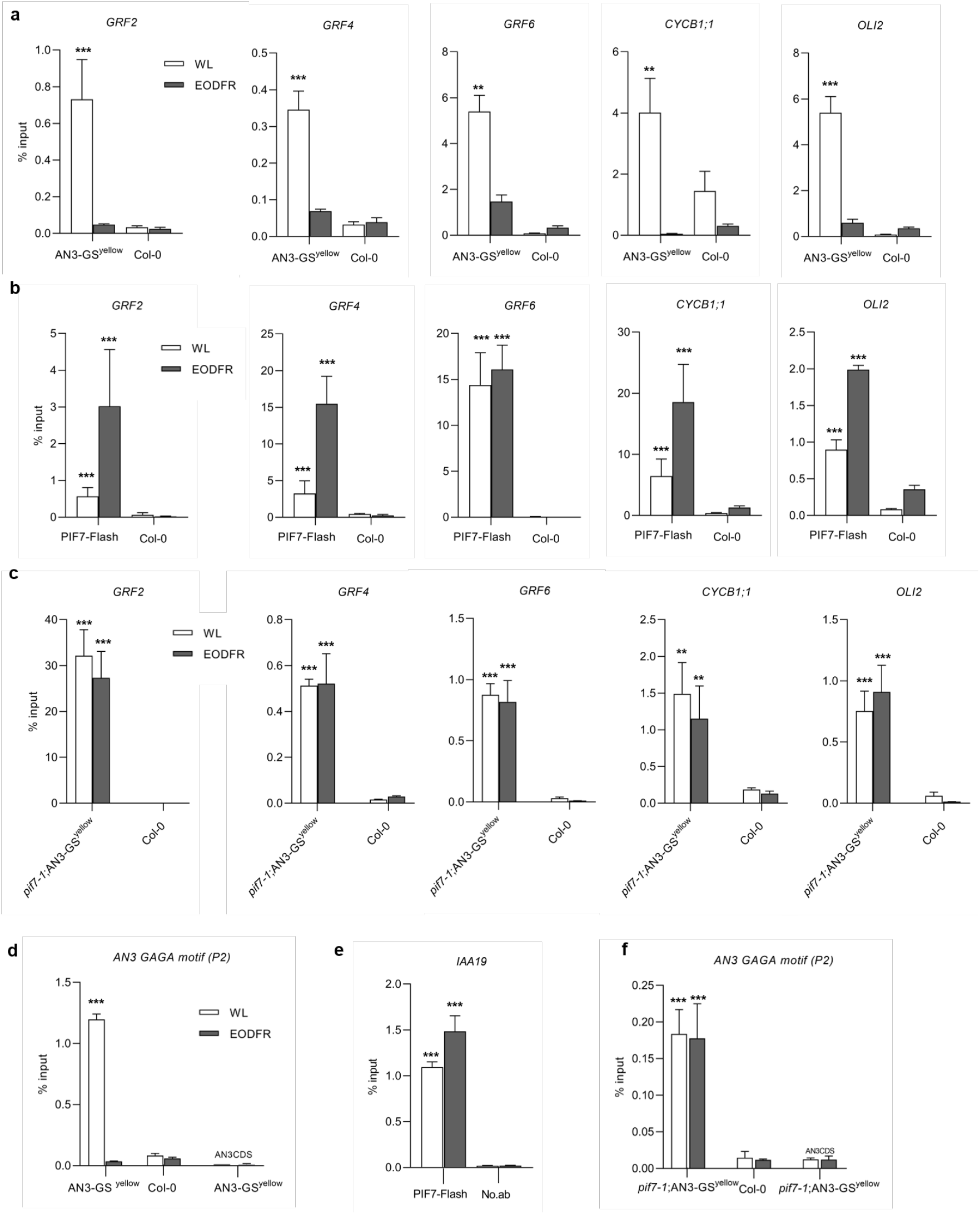
AN3 association to target gene promoters is PIF7 dependent in EODFR conditions. **a, b, c, d, e, f** ChIP assay showing enrichment of 35S::AN3-GS^yellow^, 35S::PIF7-Flash and *pif7-1*;35S::AN3-GS^yellow^ on *GRF2, GRF4, GRF6, CYCB1;1, OLI2,* and *IAA19* promoter fragments containing the G-box or PBE motif. Seedlings were grown under a light : dark (LD) 12 h : 12 h photoperiod, at 22 °C. On day 13, seedlings were treated with EODFR or kept in white light control condition. Samples were taken at ZT14 two hours after the EODFR treatment. The 35S::AN3-GS^yellow^*, pif7-1;*35S::AN3-GS^yellow^ and 35S::PIF7-Flash were incubated with an anti-GFP and anti-Myc antibodies and precipitated by Dynabeads protein A. The enrichment of fragments was determined by qPCR. The wild type (Col-0) plants acted as a control. Error bars represents the s.d. of three biological replicates. The center of the error bars represents the mean values. (\**p* < 0.05, \*\**p* < 0.01 and \*\*\**p* < 0.001, Student’s *t*-test).

## Discussion

SAS is an important survival strategy that enables plants to adapt to and thrive in often light depleted and FR-rich vegetation environments. A prominent feature of SAS is the marked reduction in leaf size, which is brought about by restricting cell proliferation and/or expansion (Carabelli et al., 2007; 2018; Patel et al., 2013; Romanowski et al., 2021). PhyB is known to play a pivotal role in orchestrating these changes in leaf development, yet current knowledge of how this is elicited is scant. This study brings a molecular level, mechanistic understanding of how phytochrome signalling is coupled to leaf blade cell division. We demonstrate that when phyB is active, cell proliferation is promoted by AN3, but following phyB deactivation by EODFR, PIF7 directly blocks AN3 action by preventing AN3 complexing at target promoters (Fig.6).

**Fig. 6:**
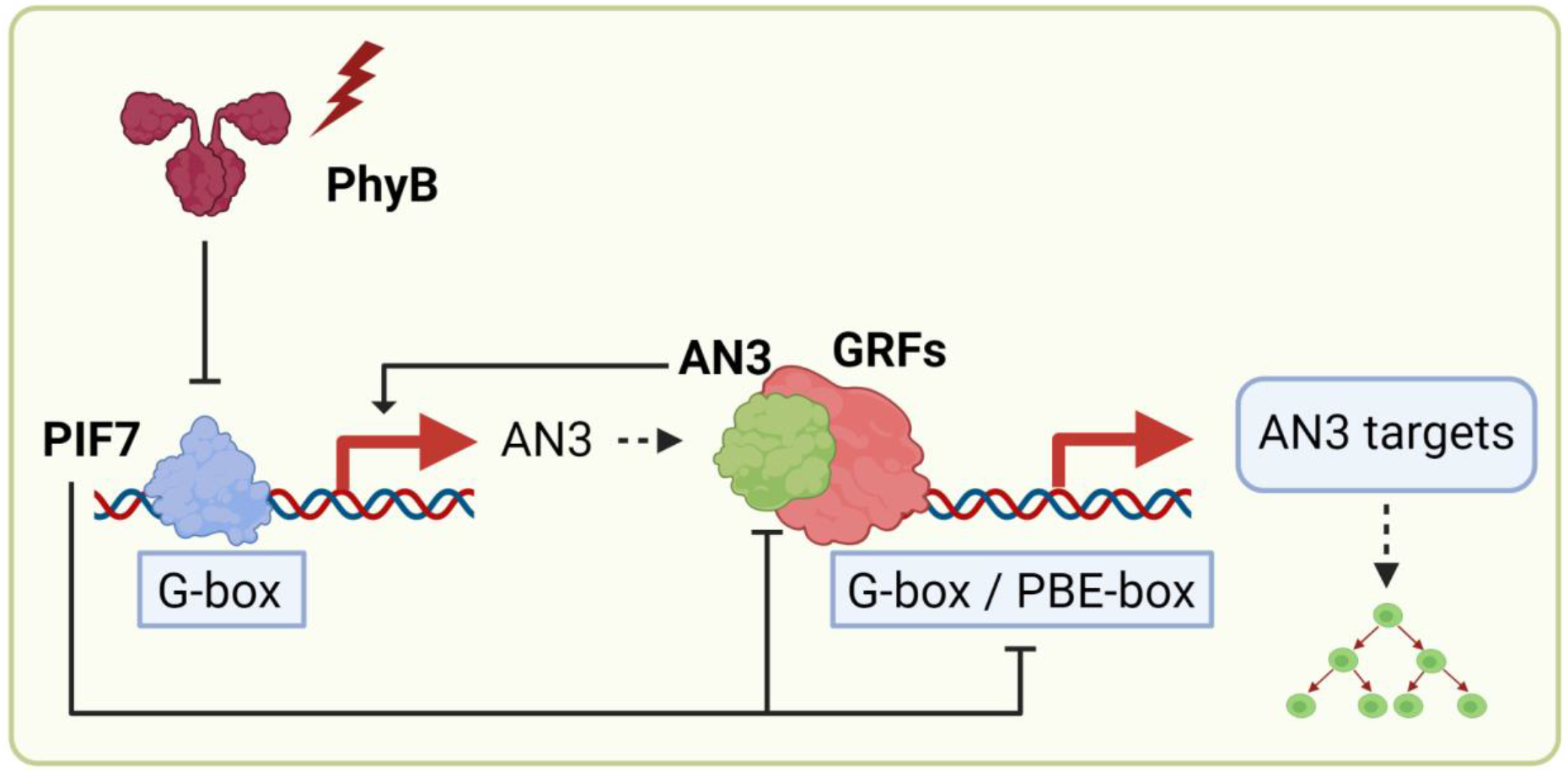
Schematic model of PIF7 control of leaf cell proliferation *via* an AN3 substitution-repression mechanism. EODFR (low R:FR ratios) inactivates phyB and increases a rapid activity of PIF7 protein. PIF7 then blocks the action of AN3 by directly binding to the promoters of multiple AN3 target genes and inhibits their expression, thereby negatively regulating leaf expansion.

Through our study we used EODFR as a tool to deactivate phytochrome, having first shown this daily treatment or the *phyB-9* allele are equally effective in repressing L3 epidermal and palisade cell proliferation (Supplementary Fig. 1a-i). By applying EODFR at different intervals through L3 development we identified two periods during leaf development (10-14- and 6-34-days post-germination) during which phyB controls cell division (Fig. 1a-h). These two proliferative phases correspond to early leaf growth, initiated at the flanks of the shoot apical meristem and the meristemoid phase of cell proliferation (Bergmann and Sack, 2007; Vercruysse et al., 2020). Supporting this interpretation, our mRNAseq data identified EODFR-controlled regulators of these two cell proliferation pathways (Supplementary Fig. 3a-c and Supplementary Fig. 10). Amongst the former group, we noted *AN3, GRF2, GRF4, GRF6 and BRM* genes that were strongly suppressed by EODFR light. As AN3 is known to operate in a complex with GRF proteins, and SWI/SNF chromatin remodelling components such as BRM, this marked the AN3 complex as a candidate for phyB regulation (Vercruyssen et al., 2014; Shimano et al., 2018; Jeong Hoe Kim. 2019).

Confirmatory evidence came from genetic analysis showing phenotypic similarities between *an3-4*, the multiple *grf1-3;grf3-1;grf5-2, grf1-3;grf3-1; grf4-1;grf5-2* mutants and *phyB-9,* all of which have small L3 blades with reduced cell number and complete insensitivity to EODFR (Fig. 2a-b and Fig. 3a-b). As GRFs are known to have functional redundancy, this data points to a potentially central role for AN3 in phyB mediated control of L3 cell division. Our data indicate AN3 promotes L3 growth in control conditions when phyB is active, but not following phyB inactivating EODFR (Fig. 2a,b and Supplementary Fig. 4b).

It is well established that FR-deactivation of phyB leads to the activation and/or accumulation of PIF transcription factors, and that PIF7 is the principal EODFR responder (Mizuno et al., 2015; Leivar et al., 2020). We, therefore, reasoned that PIF7 is required to repress cell division under EODFR. Indeed, we found that *pif7-1*, but not the *pifq* mutant, completely lacked the EODFR reduction in leaf size and epidermal cell number observed in WT plants (Fig 3a-b and Supplementary Fig. 5). This data illustrates that PIF7 suppresses cell proliferation and leaf growth following the deactivation of phytochrome with EODFR. Further, *pif7-1;an3-4* double mutant analysis showed complete *an3-4* epistasis over *pif7-1* under EODFR, indicating that PIF7 likely operates by repressing AN3 action.

It is worth noting that concurring with previously published reports (Horiguchi et al., 2005; Kawade et al., 2013), we also observed an increase in cell size in *an3-4* plants but showed that this response is suppressed by EODFR when PIF7 is present (Fig. 3c). This appears to be a PIF7 dependent compensatory mechanism to prevent leaf growth through cell expansion that would otherwise occur when AN3 is deactivated.

As EODFR led to a substantial reduction in *AN3* transcript levels, this suggested PIF7 may suppress *AN3* gene expression. This proposition was substantiated by genetic data, while ChIP-qPCR assays, showed PIF7 can directly bind to a G-box containing region of the *AN3* promoter (Fig. 4b). Collectively our data suggest that when activated by EODFR, PIF7 suppresses AN3 transcription through direct association with the *AN3* promoter.

We next reasoned that if PIF7 acted solely through the transcriptional repression of *AN3*, then AN3 overexpression should block, or at least reduce, the impact of EODFR. However, 35S::AN3-GS^yellow^ plants displayed a WT EODFR response (Supplementary Fig. 7b), rejecting this hypothesis and suggesting that PIF7 may also control AN3 through an alternative, post-transcriptional mechanism. To pursue this, we tested whether EODFR altered the ability of AN3 to bind to the UTR-located GAGA-motif and the CATGTT-motif its own promoter (Vercruyssen et al., 2014; Meng et al., 2018). Our ChIP-qPCR assays showed AN3-GS^yellow^ enrichment at these locations in control but not in EODFR (Fig. 4e-f). Further, while we observed PIF7-Flash enrichment at the G-Box region of the known target gene *IAA19* (Fig. 4d), as expected, PIF7-Flash did not bind to *AN3* GAGA and CATGTT regulatory regions in control or EODFR conditions (Fig. 4e-f). These results indicated that PIF7 may operate by removing AN3 from self-regulatory motifs in its own promoter. Concurring with this interpretation, our *in vivo* co-IP assays illustrate PIF7-Flash can directly bind AN3-GS^yellow^ indicating that PIF7 may indeed repress AN3 action through a sequestration-type mechanism (Fig. 4h).

This functional property of PIF7 is reminiscent of mechanisms reported for HFR1, PAR1/2, HEC1/2 and DELLAs, (Li et al., 2016; Hornitschek et al., 2009; Hao et al., 2012; Zhu et al., 2016). HFR1, PAR1/2, and HEC1/2 are HLH proteins that, unlike PIF7, do not bind to DNA directly but have been shown to sequester DNA-binding PIFs through heterodimerization (Hornitschek et al., 2009; Hao et al., 2012; Shi et al., 2013; Zhu et al., 2016). Interestingly, we recorded increased AN3 protein levels following EODFR (Fig. 4h). While we do not know if this results from the PIF7-AN3 interaction, sequestration has been shown to alter target protein stability. For example, DELLAs inhibit PIF binding to target genes by directly sequestering their DNA-recognition domains and by inducing PIF degradation through the ubiquitin-proteasome system (Li et al., 2016). Similarly, HFR1 can sequester and promote the degradation of PIF1 and PIF5 in the dark in a heterodimerization-dependent manner (Xu et al., 2017). PIF7 sequestration of AN3 potentially leads to the build-up of inactive heterodimers.

Our genetic data provide evidence that several key cell proliferation genes (*GRF1, GRF2, GRF3, GRF4, GRF5, GRF6, CYCB1;1, OLI2, STZ, CDC45, CDC6,* and *DEL1*) are antagonistically regulated by AN3 and PIF7. As the known PIF7 and AN3 binding G-box/PBE-boxes, were common motifs found in the promoters of all of these genes, this suggested PIF7 and AN3 signal convergence at common motifs (Vercruyssen et al., 2014; Galvao et al., 2019). ChIP-qPCR assays showed that this was indeed the case and that EODFR induced switching from AN3 to PIF7 enrichment. The loss of AN3-eviction in the *pif7-1* mutant confirmed that AN3 binding is dependent on the absence of active PIF7 (Fig. 5c and Supplementary Fig. 8c). The opposing action of AN3 and PIF7 on target gene expression means that the EODFR-induced substitution mechanism facilitates the on to off modulation of gene expression.

Interestingly, analogous behaviour has previously been reported for AN3, which can competitively interact with JANUS to inhibit its Pol II recruitment on PLT1 promoter and antagonistically regulate PLT1 transcription in the root meristem (Xiong et al., 2020). Further, opposing action has been observed for PIF1/3 and the bZIP transcription factors, HY5 and HYH which can physically interact and can bind to G-box containing regions in a common set of target promoters (Chen et al., 2013; Toledo-Ortiz et al., 2014). Our findings also have similarities to the recently reported PIF3-TCP4 substitution-suppression module (Dong et al., 2019). In this instance, darkness induced accumulation of PIF3 in seedling cotyledons triggers the substitution of TCP4 for PIF3 at shared cis-elements in *SAUR* gene promoters (Dong et al., 2019). As for the PIF7-AN3 module, conditional switching between PIF1/3 and HY5/HYH or PIF3 and TCP4 promoter occupancy provides a mechanism through which external signals can direct changes in gene expression.

Previously PIF7 has been shown to participate in transcriptional activation of key SAS genes including *YUC8, YUC9, IAA19, IAA29, ATHB2 and PRE1* (de Wit et al., 2015; Mizuno et al., 2015; Peng et al., 2018). Further, PIF7 can directly interact with MORF RELATED GENE 2 (MRG2), to promote histone acetylation and the expression of *YUC8, IAA19* and *PRE1*. In this study, we have shown PIF7 acts as a transcriptional repressor, as it does in the regulation of the DRE-Binding1 C-repeat binding factor (DREB1C) (Kidokoro et al., 2009). Interestingly, PIF3 and PIF1 have both been reported to interact with HISTONE DEACETYLASE 15 (HDA15) to repress gene expression (Liu et al., 2013; Gu et al., 2017). The mode of action through which PIF7 deactivates AN3 and suppresses target gene expression also has similarities to the well-studied mammalian *hairy and enhancer of split-1/ Hairy/enhancer-of-split related with YRPW motif protein* (*hes/hey*) system. Hes and Hes are bHLH notch signalling proteins, closely related to the Drosophila *hairy/enhancer of split* family of genes (Weber et al., 2014). Like PIFs, Hes and Hey proteins preferentially bind DNA at E-Box (CANNTG) variants including the G-Box. Studies indicate that Hes and Hey mainly act as co-repressors by interacting with and suppressing the activity of transcription factors, and through the recruitment of deacetylases (Iso et al., 2001; Takata & Ishikawa, 2003; Gould et al., 2009, Weber et al., 2014). It will be interesting to establish whether the PIF7-mediated AN3 promoter eviction and transcriptional suppression represent an analogous process that is conserved across species.

In conclusion, this study shows PIF7 inhibits leaf cell proliferation by inactivating the central leaf developmental regulator AN3. EODFR-activated PIF7 represses *AN3* expression by directly interacting with and sequestering AN3 from cis-elements in its own promoter. We show PIF7 and AN3 signalling converges at G-box/PBE-box promoter elements in a major set of genes that control cell division. EODFR treatment induces the substitution of AN3 for PIF7 at target promoters and vitally, a switch from promotion to repression of cell cycle gene expression.

## Methods

### Plant material and growth conditions

All *Arabidopsis thaliana* mutants and transgenic plants that were used in this study were from the Columbia (Col-0) ecotype. Some of the mutant and overexpressing lines used in this study were described elsewhere including PIF7-Flash (35S::PIF7-Flash; 9xMyc-6xHis-3xFlag) (Li et al. 2012), 35S::AN3-GS^yellow^ (Besbrugge et al., 2018). The triple mutant *grf1-3;grf3-1;grf5-2,* and quintuple mutant *grf1-3;grf3-1;grf4-1;grf5-2* have been previously described elsewhere (Lee et al., 2018). The *phyB-9* (Reed et al. 1993), *pif7-1* (Leivar et al. 2008a), *pif1-2, pif3-3, pif4-2, pif5-2* (*pifq*) (Leivar et al. 2008b) have been previously characterized. *an3-4* is an AN3 null mutant derived from an X-ray-irradiated population of the Col-0 accession (Horiguchi et al., 2005). New mutant combinations included *an3-4;phyB-9*, which were obtained by crossing *an3-4* with *phyB-9* plants, and *an3-4;pif7-1* were obtained by crossing *an3-4* with *pif7-1* plants and the triple *an3-4;phyB-9;pif7-1* mutant were obtained by crossing *an3-4;phyB-9* with *an3-4;pif7-1* plants. 35S::PIF7-Flash; 35S::AN3-GS^yellow^ plants were obtained by crossing 35S::PIF7-Flash with 35S::AN3-GS^yellow^ plants and double homozygous transgenic plants were screened out with 30 μg ml^-1^ hygromycin and 50 μg ml^-1^ kanamycin. *pif7-1;*35S::AN3-GS^yellow^ plants were obtained by crossing 35S::AN3-GS^yellow^ with *pif7-1* plants. *an3-4* and *pif7-1* mutations were identified by PCR. The *phyB-9* mutation was confirmed by sanger sequencing. The absence of the secondary *VENOSA4* mutation (Yoshida et al., 2018), was confirmed by sequencing. All primers used in this work are listed in Supplementary Table 2.

Seeds were sown on F2 + S Levington Advance Seed and Modular Compost plus Sand soil mix (ICL Specialty Fertilizers, Suffolk, U.K.) and stratified for four days in darkness at 4 ºC. Seeds were grown inside a Percival SE-41L cabinet (CLF Plant Climatics, Wertingen, Germany) under a light: dark (LD) 12 h : 12 h photoperiod, at 100 μmol m^−2^ s^−1^ fluence rate and 22 °C of constant temperature. Half of the plants exposed to daily EODFR (40 μmol m^−2^ s^−1^) from Day 6 (before L3 emergence) until sampling on Day 34. For EODFR treatments, we used seven 24V OSLON 150 ILS-OW06-FRED-SD111 FR led strips (Intelligent Led Solutions, Berkshire, UK), to deliver 40 µmol m^−2^ s^−1^ of FR light (730 nm) for 10 min. The spectrum of both light sources can be found in Supplementary Fig. 11. All reagents used in this work were purchased from Merck KGaA (Darmstadt, Germany) unless otherwise specified. Further growth conditions details are provided in the respective figure legends.

### Blade area measurement

Whole leaf pictures for blade area measurements were taken from a camera stand using a Nikon G20 camera with automatic focus settings (Romanowski et al., 2021). A ruler was included in each photograph for scaling purposes. Blade area was measured using NIH ImageJ software (http://rsb.info.nih.gov/nih-image/). Bar charts and box plots were generated using Prism 8 (GraphPad Software, San Diego, CA).

### Generation of transparent leaf blades for microscopy imaging

Leaf blades were excised from 34 D.A.S plants with a razor blade and cleared as described in (Romanowski et al., 2021). Blades were then mounted onto microscope slides with the adaxial layer facing down.

### Cell size and number measurement

For epidermal and palisade cell parameter determination cleared blades (34 D.A.S) were mounted and visualized using an Eclipse E600 (Nikon) DIC microscope using either a 10X or a 20X objective. Individual abaxial epidermal and adaxial subepidermal palisade cell sizes were measured using NIH ImageJ software (http://rsb.info.nih.gov/nih-image/). Average leaf cell sizes were obtained by deriving the mean values of 9 adjacent cells from the base, middle, and tip sections of each leaf, or these sections combined. The average total number of cells was obtained by dividing the blade area by the total cell size of each blade, and then averaging the mean total number of cells of each blade. Average cell density was obtained by dividing the total number of cells by the blade area and then averaging the mean cell density of each blade (Romanowski et al., 2021). An S8 stage mic 1 mm/0.01 mm DIV graticule (#02A00404, Pyser-SGI Ltd., Kent, UK) was used for scaling. Bar charts and box plots were generated using Prism 8 (GraphPad Software, San Diego, CA).

### RNA extraction and RT-qPCR

For gene expression analysis, plants were grown for 13 days under a light : dark (LD) 12 h : 12 h photoperiod, at 100 μmoles m^−2^ s^−1^ fluence rate and 22 °C of constant temperature. On day 13, plants were either shifted to EODFR of 40 μmoles m^−2^ s^−1^ FR light or kept in 100 μmoles m^−2^ s^−1^ white light condition. Briefly, 13 days old whole seedlings were harvested at zeitgeber 22 (ZT) in RNA later solution (Thermofisher) and leaf 3 blades were dissected with a razor, in a Petri dish filled with RNA later solution, under a Leica MZ 16 F dissecting microscope. Total RNA was extracted using the RNeasy Plant Mini Kit (Qiagen) with on-column DNase digestion. All samples were processed on the same day. cDNA synthesis was performed using the qScript cDNA SuperMix (Quanta Biosciences) as described by the manufacturer. The RT-qPCR was set up as a 10 μL reaction using Lightcycler 480 SYBR Green Master Mix (Roche) in a 384-well plate, performed with a Lightcycler 480 system (Roche). Results were analyzed using the Light Cycler 480 software. Gene-specific primers are listed in Supplementary Table 2. Bar charts were generated using Prism 8 (GraphPad Software, San Diego, CA).

### ChIP-qPCR assay

For ChIP-qPCR assay, plants were grown for 13 days under a light : dark (LD) 12 h : 12 h photoperiod, at 100 μmoles m^−2^ s^−1^ fluence rate and 22 °C of constant temperature. On day 13, plants were either shifted to EODFR of 40 μmoles m^−2^ s^−1^ FR light or kept in 100 μmoles m^−2^ s^−1^ white light condition. In all genotypes and treatments, 13 day old whole seedling shoots were harvested at ZT14, 2 h after the EODFR pulse. Chromatin immunoprecipitation analyses protocol was adapted from (Gendrel et al., 2002). Briefly, whole above ground seedlings were fixed under vacuum for 15 min repeated twice at 25 PSI in 1X PBS containing 1% formaldehyde. The reaction was quenched by adding 2 M glycine to a final concentration of 125 mM and incubated for 5 min. Samples were washed three times in sterile MilliQ H_2_O and ground to a fine powder with liquid nitrogen. Nuclei were isolated in three steps with the series of extraction buffers EB1-3. The lysate was centrifugated at 4,000 rpm at 4 °C for 20 min in EB1 (0.4 M sucrose, 10 mM Tris-HCl (pH8), 10 mM MgCl_2_, 5 mM 2-mercaptoethanol (BME), 0.2 mM phenylmethylsulfonyl fluoride (PMSF), 2 Phosphatase Inhibitor Mini Tablets, 1 mM EDTA), and then at 12,000 rpm for 10 min in EB2 (0.25 M sucrose, 10 mM Tris-HCl (pH 8), 10 mM MgCl_2_, 5 mM BME, 0.2 mM PMSF, 1 complete mini tablet, 1 mM EDTA) and then at 15,000 rpm for 1h in EB3 (1.7 M sucrose, 10 mM Tris-HCL (pH 8), 2 mM MgCl_2_, 5 mM BME, 0.2 mM PMSF, 0.15% (v/v) Triton X-100, 1 complete mini tablet, 1 mM EDTA). Chromatin was extracted with cold nuclei lysis buffer (50 mM Tris-HCl (pH 8), 10 mM EDTA, 0.4 mM PMSF, 1 complete mini tablet,1% w/v SDS) after centrifugation at 4000 rpm for 5 min. The chromatin solution was sonicated for 7 x 10s, 5 x 10s and 3 x 10s at power setting 9. The chromatin solution was diluted to dilute the 1% SDS to 0.1% SDS with ChIP buffer (16.7 mM Tris-HCL (pH 8), 167 mM NaCl, 0.2 mM PMSF, 1.1% (v/v) Triton X-100, 2.5 complete mini tablet, 1.2 mM EDTA). The chromatin complexes were immunoprecipitated with anti-GFP antibody (ab290) and an anti-myc mouse antibody (mAb 9E10, Calbiochem) with a concentration of 2 μg / sample and incubated in ChIP dilution buffer with 20 μL of Dynabeads Protein A (Thermofisher) overnight at 4°C. An equal amount of chromatin solution was not treated with antibody and, thus, served as the mock antibody control. The beads were washed for 5 min each time at 4 °C with 1 ml of each of the following buffers: 2 times with low salt wash buffer (150 mM NaCl, 20 mM Tris-HCl (pH 8), 0.1% w/v SDS, 1% v/v Triton X-100, and 2 mM EDTA), two times with high salt wash buffer (500 mM NaCl, 20 mM Tris-HCl (pH 8), 0.1% w/v SDS, 1% v/v Triton X-100, and 2 mM EDTA), two times with LiCl wash buffer (0.25 M LiCl, 1% w/v sodium deoxycholate, 10 mM Tris-HCl (pH 8), 1% v/v NP-40, and 1 mM EDTA), and two times with TE buffer (1 mM EDTA and 10 mM Tris-HCl pH 8). DNA was extracted from the beads with elution buffer containing 1% w/v SDS and 0.1 M NaHCO_3_ at 65 °C for 15 min and reversely cross-linked with 192 mM NaCl at 65 °C overnight. Proteins were removed with an equal volume of phenol:chloroform:isoamyl-alcohol (25:24:1). An additional step was performed by incubating the chromatin solution for 1h at 45°C in a buffer solution containing 0.5 M EDTA, 1M Tris-HCl (pH6.5) and 10 mg/mL proteinase K to elute and remove any of the remaining proteins. DNA was precipitated with 2.5 volume of 100% ethanol, 1 μl glycogen, 1/10 volume of 1.5 M potassium acetate (pH 5.2) and spun at 15,000 rpm for 30 min. The supernatant was removed, and the DNA pellet was dried with 70% ethanol. A small aliquot of the untreated sonicated chromatin was reverse cross-linked for use as the total input DNA control. ChIP assays were quantified by qPCR after normalising with the input DNA. Gene-specific primers covering the G-box and PBE-box are listed in Supplementary Table 2. Bar charts were generated using Prism 8 (GraphPad Software, San Diego, CA).

### Co-IP assays

For co-immunoprecipitation assays, plants were grown for 13 days under a light : dark (LD) 12 h : 12 h photoperiod, at 100 μmoles m^−2^ s^−1^ fluence rate and 22 °C of constant temperature. On day 13, plants were either shifted to EODFR of 40 μmoles m^−2^ s^−1^ FR light or kept in 100 μmoles m^−2^ s^−1^ white light condition. The whole seedling shoots of 35S::PIF7-Flash;35S::AN3-GS^yellow^ double overexpressor and 35S::AN3-GS^yellow^ single overexpressor were harvested in liquid nitrogen at ZT14. The Co-IP was performed following the method of Zhu et al., 2018. Briefly, samples were ground to a fine powder in liquid nitrogen and homogenized in two volumes (mg/μL) of Co-IP buffer containing (100 mM phosphate buffer, pH 7.8, 150 mM NaCl, 0.1% NP40, 1X protease inhibitor cocktail, 1 mM PMSF, 10 mM iodoacetamide, 40 μM bortezomib, 25 mM β-glycerophosphate, 10 mM sodium fluoride (NaF), and 2 mM sodium orthovanadate (Na_3_VO_4_)). After centrifugation at 15,000 rpm for 15 min at 4 °C in the dark, total protein levels were quantified with the Bradford Protein Assay (Bio-Rad, USA) and the lysate was incubated with anti-Myc antibody pre coupled Dynabeads Protein A (Thermofisher) for 4 h at 4 °C. The samples were precipitated with anti-Myc antibody and ran on a 4-12% NuPAGE Bis-Tris gel (ThermoFisher) and blotted using the MiniBlot Module (Life Technologies) at 20V for 1 hour. Western blot detection for PIF7-Flash was performed using anti-Myc 9B11 (Cell Signalling) 1:1000, and anti-Mouse IgG (whole molecule)-Peroxidase antibody (A4416 Sigma) 1:10000. Western blot detection for AN3-GS^yellow^ was performed using an anti-GFP antibody (ab290).1:1000, and Goat Anti-Rabbit IgG H&L (HRP) (ab6721) 1:10000.

### Promoter analysis

A 2-kb upstream promoter region of each target gene was obtained using PlantPAN 3.0 (http://plantpan.itps.ncku.edu.tw/). Regions were manually searched for G-box (CACGTG) and PBE-box (CA[TG/CA]TG target binding sites.

### Statistical information

Statistical difference of two populations was tested by two-tailed, unpaired Student’s t-test. To compare three or more populations, a one-way analysis of variance (ANOVA) followed by Tukey’s test (comparison among all groups) was performed. When Tukey’s test was employed, letters were used to indicate which treatment groups were significantly different. All analyses were done using GraphPad Prism 8.0.2 (GraphPad Software) or Minitab 19 (Minitab Ltd.) unless otherwise indicated.

### Data availability

The raw numerical data that support the findings of this study are available from the corresponding author upon reasonable request. All other data supporting the findings of this study are available within the paper and its supplementary information files.

## Acknowledgements

We would like to thank Prof. Dirk Inzé for providing us with the *35S::AN3-GS*^*yellow*^ line, Prof. Gorou Horiguchi for *an3-4* seeds, Prof. Jeong Hoe Kim for *grf1-3;grf3-1;grf5-2*, *grf1-3;grf3-1; grf4-1;grf5-2* seeds and Prof. Joanne Chory and Dr Elena Monte for 35S::PIF7-Flash line. The authors would also like to thank Prof. Enamul Huq, Dr Beatriz Orosa, and Dr Gabriela Toledo-Ortiz and Dr Michael Skelly for assistance with co-IP and ChIP-qPCR assays, respectively. E.H. was supported by the Punjab Educational Endowment Fund PEEF/CMMS/2016/203. This work was supported by Biotechnology and Biological Sciences Research Council-United Kingdom Research and Innovation (BBSRCUKRI) grants BB/M025551/1 and BB/N005147/1, awarded to KJH.

## Contributions

Experimental design by E.H., A.R. and K.J.H.; Experimental work and data analysis by E.H. and A.R.; Original manuscript draft by E.H and K.J.H.; Review & Editing by E.H., A.R. and K.J.H.

## Corresponding authors

Correspondence to Karen J. Halliday.

## Supplementary information

The submitted version contains supplementary materials and is submitted as a separate PDF file.

## References

1. Sessa G, Carabelli M, Possenti M, Morelli G, Ruberti I. Multiple Pathways in the Control of the Shade Avoidance Response. Plants (Basel). 2018 Nov 17;7(4):102. doi: 10.3390/plants7040102. PMID: 30453622; PMCID: PMC6313891.c72

2. Yang D, Seaton DD, Krahmer J, Halliday KJ. Photoreceptor effects on plant biomass, resource allocation, and metabolic state. Proc Natl Acad Sci U S A. 2016 Jul 5;113(27):7667–72. doi: 10.1073/pnas.1601309113. Epub 2016 Jun 21. PMID: 27330114; PMCID: PMC4941476.

3. Krahmer J, Abbas A, Mengin V, Ishihara H, Romanowski A, Furniss JJ, Moraes TA, Krohn N, Annunziata MG, Feil R, Alseekh S, Obata T, Fernie AR, Stitt M, Halliday KJ. Phytochromes control metabolic flux, and their action at the seedling stage determines adult plant biomass. J Exp Bot. 2021 Feb 5:erab038. doi: 10.1093/jxb/erab038. Epub ahead of print. PMID: 33544130.

4. Patel D, Basu M, Hayes S, Majlath I, Hetherington FM, Tschaplinski TJ, Franklin KA (2013) Temperature-dependent shade avoidance involves the receptor-like kinase ERECTA. Plant J 73: 980–992

5. Romanowski A, Furniss JJ, Hussain E, Halliday KJ. Phytochrome regulates cellular response plasticity and the basic molecular machinery of leaf development. Plant Physiol. 2021 Mar 9:kiab112. doi: 10.1093/plphys/kiab112. Epub ahead of print. PMID: 33693822.

6. Franklin KA, Quail PH. Phytochrome functions in Arabidopsis development. J Exp Bot. 2010;61(1):11–24. doi: 10.1093/jxb/erp304. PMID: 19815685; PMCID: PMC2800801.

7. Klose C, Nagy F, Schäfer E. Thermal Reversion of Plant Phytochromes. Mol Plant. 2020 Mar 2;13(3):386–397. doi: 10.1016/j.molp.2019.12.004. Epub 2019 Dec 6. PMID: 31812690.

8. Franklin KA, Whitelam GC. Light signals, phytochromes and cross-talk with other environmental cues. J Exp Bot. 2004 Jan;55(395):271–6. doi: 10.1093/jxb/erh026. Epub 2003 Dec 12. PMID: 14673030.

9. M. J. Yanovsky J. J. Casal G. C. Whitelam. Phytochrome A, phytochrome B and HY4 are involved in hypocotyl growth responses to natural radiation in Arabidopsis: weak de‐etiolation of the phyA mutant under dense canopies. Plant, Celt and Environment (1995) 18, 788–7.

10. Park E, Park J, Kim J, Nagatani A, Lagarias JC, Choi G. Phytochrome B inhibits binding of phytochrome-interacting factors to their target promoters. Plant J. 2012 Nov;72(4):537–46. doi: 10.1111/j.1365-313X.2012.05114.x. Epub 2012 Sep 25. PMID: 22849408; PMCID: PMC3489987.

11. Park E, Kim Y, Choi G. Phytochrome B Requires PIF Degradation and Sequestration to Induce Light Responses across a Wide Range of Light Conditions. Plant Cell. 2018 Jun;30(6):1277–1292. doi: 10.1105/tpc.17.00913. Epub 2018 May 15. PMID: 29764986; PMCID: PMC6048787.

12. Pham VN, Kathare PK, Huq E. Phytochromes and Phytochrome Interacting Factors. Plant Physiol. 2018 Feb;176(2):1025–1038. doi: 10.1104/pp.17.01384. Epub 2017 Nov 14. PMID: 29138351; PMCID: PMC5813575.

13. Huang X, Zhang Q, Jiang Y, Yang C, Wang Q, Li L. Shade-induced nuclear localization of PIF7 is regulated by phosphorylation and 14-3-3 proteins in Arabidopsis. Elife. 2018 Jun 21;7:e31636. doi: 10.7554/eLife.31636. PMID: 29926790; PMCID: PMC6037483.

14. Salter MG, Franklin KA, Whitelam GC (2003) Gating of the rapid shade-avoidance response by the circadian clock in plants. Nature 426: 680–683.

15. Mizuno T, Oka H, Yoshimura F, Ishida K, Yamashino T (2015) Insight into the mechanism of end-of-day far-red light (EODFR)-induced shade avoidance responses in Arabidopsis thaliana. Biosci Biotechnol Biochem 79: 1987–1994.

16. Franklin KA. Shade avoidance. New Phytol. 2008;179(4):930–944. doi: 10.1111/j.1469-8137.2008.02507.x. Epub 2008 Jun 5. PMID: 18537892.

17. Strasser B, Sanchez-Lamas M, Yanovsky MJ, Casal JJ, Cerdan PD (2010) Arabidopsis thaliana life without phytochromes. Proc Natl Acad Sci U S A 107: 4776–4781.

18. Leivar P, Martín G, Soy J, Dalton-Roesler J, Quail PH, Monte E. Phytochrome-imposed inhibition of PIF7 activity shapes photoperiodic growth in Arabidopsis together with PIF1, 3, 4 and 5. Physiol Plant. 2020 Jul;169(3):452–466. doi: 10.1111/ppl.13123. PMID: 32412656.

19. Li L, Ljung K, Breton G, Schmitz RJ, Pruneda-Paz J, Cowing-Zitron C, Cole BJ, Ivans LJ, Pedmale UV, Jung HS, Ecker JR, Kay SA, Chory J (2012) Linking photoreceptor excitation to changes in plant architecture. Genes Dev 26: 785–790.

20. Jiang Y, Yang C, Huang S, Xie F, Xu Y, Liu C, Li L. The ELF3-PIF7 Interaction Mediates the Circadian Gating of the Shade Response in Arabidopsis. iScience. 2019 Dec 20;22:288–298. doi: 10.1016/j.isci.2019.11.029. Epub 2019 Nov 20. PMID: 31805433; PMCID: PMC6909221.

21. Carabelli M, Possenti M, Sessa G, Ciolfi A, Sassi M, Morelli G, Ruberti I (2007) Canopy shade causes a rapid and transient arrest in leaf development through auxin-induced cytokinin oxidase activity. Genes Dev 21: 1863–1868

22. Carabelli M, Possenti M, Sessa G, Ruzza V, Morelli G, Ruberti I (2018) Arabidopsis HD-Zip II proteins regulate the exit from proliferation during leaf development in canopy shade. J Exp Bot 69: 5419–5431

23. Tsukaya H, Kozuka T, Kim GT. Genetic control of petiole length in Arabidopsis thaliana. Plant Cell Physiol. 2002 Oct;43(10):1221–8. doi: 10.1093/pcp/pcf147. PMID: 12407202.

24. Kim JH, Kende H. 2004. A transcriptional coactivator, AtGIF1, is involved in regulating leaf growth and morphology in Arabidopsis. Proceedings of the National Academy of Sciences, USA 101, 13374–13379.

25. Horiguchi G, Kim GT, Tsukaya H. The transcription factor AtGRF5 and the transcription coactivator AN3 regulate cell proliferation in leaf primordia of Arabidopsis thaliana. Plant J. 2005 Jul;43(1):68–78. doi: 10.1111/j.1365-313X.2005.02429.x. PMID: 15960617.

26. Vercruyssen L, Verkest A, Gonzalez N, Heyndrickx KS, Eeckhout D, Han SK, Jégu T, Archacki R, Van Leene J, Andriankaja M, De Bodt S, Abeel T, Coppens F, Dhondt S, De Milde L, Vermeersch M, Maleux K, Gevaert K, Jerzmanowski A, Benhamed M, Wagner D, Vandepoele K, De Jaeger G, Inzé D. ANGUSTIFOLIA3 binds to SWI/SNF chromatin remodeling complexes to regulate transcription during Arabidopsis leaf development. Plant Cell. 2014 Jan;26(1):210–29. doi: 10.1105/tpc.113.115907. Epub 2014 Jan 17. PMID: 24443518; PMCID: PMC3963571

27. Nelissen H, Eeckhout D, Demuynck K, Persiau G, Walton A, van Bel M, Vervoort M, Candaele J, De Block J, Aesaert S, Van Lijsebettens M, Goormachtig S, Vandepoele K, Van Leene J, Muszynski M, Gevaert K, Inzé D, De Jaeger G. Dynamic Changes in ANGUSTIFOLIA3 Complex Composition Reveal a Growth Regulatory Mechanism in the Maize Leaf. Plant Cell. 2015 Jun;27(6):1605–19. doi: 10.1105/tpc.15.00269. Epub 2015 Jun 2. PMID: 26036253; PMCID: PMC4498210.

28. Besbrugge N, Van Leene J, Eeckhout D, Cannoot B, Kulkarni SR, De Winne N, Persiau G, Van De Slijke E, Bontinck M, Aesaert S, Impens F, Gevaert K, Van Damme D, Van Lijsebettens M, Inzé D, Vandepoele K, Nelissen H, De Jaeger G. GSyellow, a Multifaceted Tag for Functional Protein Analysis in Monocot and Dicot Plants. Plant Physiol. 2018 Jun;177(2):447–464. doi: 10.1104/pp.18.00175. Epub 2018 Apr 20. PMID: 29678859; PMCID: PMC6001315.

29. Shimano S, Hibara KI, Furuya T, Arimura SI, Tsukaya H, Itoh JI. Conserved functional control, but distinct regulation, of cell proliferation in rice and Arabidopsis leaves revealed by comparative analysis of GRF-INTERACTING FACTOR 1 orthologs. Development. 2018 Apr 5;145(7):dev159624. doi: 10.1242/dev.159624. PMID: 29567670.

30. Kozuka T, Horiguchi G, Kim GT, Ohgishi M, Sakai T, Tsukaya H. The different growth responses of the Arabidopsis thaliana leaf blade and the petiole during shade avoidance are regulated by photoreceptors and sugar. Plant Cell Physiol. 2005 Jan;46(1):213–23. doi: 10.1093/pcp/pci016. Epub 2005 Jan 19. PMID: 15659441.

31. Andriankaja M, Dhondt S, De Bodt S, Vanhaeren H, Coppens F, De Milde L, Mühlenbock P, Skirycz A, Gonzalez N, Beemster GT, Inzé D. Exit from proliferation during leaf development in Arabidopsis thaliana: a not-so-gradual process. Dev Cell. 2012 Jan 17;22(1):64–78. doi: 10.1016/j.devcel.2011.11.011. Epub 2012 Jan 5. PMID: 22227310.

32. Beltramino M, Ercoli MF, Debernardi JM, Goldy C, Rojas AML, Nota F, Alvarez ME, Vercruyssen L, Inzé D, Palatnik JF, Rodriguez RE. Robust increase of leaf size by Arabidopsis thaliana GRF3-like transcription factors under different growth conditions. Sci Rep. 2018 Sep 7;8(1):13447. doi: 10.1038/s41598-018-29859-9. PMID: 30194309; PMCID: PMC6128883.

33. Gonzalez N, Vanhaeren H, Inzé D. Leaf size control: complex coordination of cell division and expansion. Trends Plant Sci. 2012 Jun;17(6):332–40. doi: 10.1016/j.tplants.2012.02.003. Epub 2012 Mar 6. PMID: 22401845.

34. Lee BH, Ko JH, Lee S, Lee Y, Pak JH, Kim JH. The Arabidopsis GRF-INTERACTING FACTOR gene family performs an overlapping function in determining organ size as well as multiple developmental properties. Plant Physiol. 2009 Oct;151(2):655–68. doi: 10.1104/pp.109.141838. Epub 2009 Jul 31. PMID: 19648231; PMCID: PMC2754652.

35. Van Leene J, Hollunder J, Eeckhout D, Persiau G, Van De Slijke E, Stals H, Van Isterdael G, Verkest A, Neirynck S, Buffel Y, De Bodt S, Maere S, Laukens K, Pharazyn A, Ferreira PC, Eloy N, Renne C, Meyer C, Faure JD, Steinbrenner J, Beynon J, Larkin JC, Van de Peer Y, Hilson P, Kuiper M, De Veylder L, Van Onckelen H, Inzé D, Witters E, De Jaeger G. Targeted interactomics reveals a complex core cell cycle machinery in Arabidopsis thaliana. Mol Syst Biol. 2010 Aug 10;6:397. doi: 10.1038/msb.2010.53. PMID: 20706207; PMCID: PMC2950081.

36. Kawade K, Horiguchi G, Usami T, Hirai MY, Tsukaya H. ANGUSTIFOLIA3 signaling coordinates proliferation between clonally distinct cells in leaves. Curr Biol. 2013 May 6;23(9):788–92. doi: 10.1016/j.cub.2013.03.044. Epub 2013 Apr 18. PMID: 23602479.

37. Liebsch D, Palatnik JF. MicroRNA miR396, GRF transcription factors and GIF co-regulators: a conserved plant growth regulatory module with potential for breeding and biotechnology. Curr Opin Plant Biol. 2020 Feb;53:31–42. doi: 10.1016/j.pbi.2019.09.008. Epub 2019 Nov 11. PMID: 31726426.

38. Kim JS, Mizoi J, Kidokoro S, Maruyama K, Nakajima J, Nakashima K, Mitsuda N, Takiguchi Y, Ohme-Takagi M, Kondou Y, Yoshizumi T, Matsui M, Shinozaki K, Yamaguchi-Shinozaki K. Arabidopsis growth-regulating factor7 functions as a transcriptional repressor of abscisic acid- and osmotic stress-responsive genes, including DREB2A. Plant Cell. 2012 Aug;24(8):3393–405. doi: 10.1105/tpc.112.100933. Epub 2012 Aug 31. PMID: 22942381; PMCID: PMC3462639.

39. Omidbakhshfard MA, Fujikura U, Olas JJ, Xue GP, Balazadeh S, Mueller-Roeber B. GROWTH-REGULATING FACTOR 9 negatively regulates arabidopsis leaf growth by controlling ORG3 and restricting cell proliferation in leaf primordia. PLoS Genet. 2018 Jul 9;14(7):e1007484. doi: 10.1371/journal.pgen.1007484. PMID: 29985961; PMCID: PMC6053248.

40. Xie Y, Liu Y, Wang H, Ma X, Wang B, Wu G, Wang H. Phytochrome-interacting factors directly suppress MIR156 expression to enhance shade-avoidance syndrome in Arabidopsis. Nat Commun. 2017 Aug 24;8(1):348. doi: 10.1038/s41467-017-00404-y. PMID: 28839125; PMCID: PMC5570905.

41. Galvāo VC, Fiorucci AS, Trevisan M, Franco-Zorilla JM, Goyal A, Schmid-Siegert E, Solano R, Fankhauser C. PIF transcription factors link a neighbor threat cue to accelerated reproduction in Arabidopsis. Nat Commun. 2019 Sep 5;10(1):4005. doi: 10.1038/s41467-019-11882-7. PMID: 31488833; PMCID: PMC6728355.

42. Liu Z, Li N, Zhang Y, Li Y. Transcriptional repression of GIF1 by the KIX-PPD-MYC repressor complex controls seed size in Arabidopsis. Nat Commun. 2020 Apr 15;11(1):1846. doi: 10.1038/s41467-020-15603-3. PMID: 32296056; PMCID: PMC7160150.

43. Peng, M., Li, Z., Zhou, N., Ma, M., Jiang, Y., Dong, A., Shen, W. H. and Li, L. (2018) ‘Linking phytochrome-interacting factor to histone modification in plant shade avoidance’, Plant Physiology. American Society of Plant Biologists, 176(2), pp. 1341–1351. doi: 10.1104/pp.17.01189.

44. Meng, L. S., Li, C., Xu, M. K., Sun, X. D., Wan, W., Cao, X. Y., Zhang, J. L. and Chen, K. M. (2018) ‘Arabidopsis ANGUSTIFOLIA3 (AN3) is associated with the promoter of CONSTITUTIVE PHOTOMORPHOGENIC1 (COP1) to regulate light-mediated stomatal development’, Plant Cell and Environment. Blackwell Publishing Ltd, 41(7), pp. 1645–1656. doi: 10.1111/pce.13212.

45. Bergmann DC, Sack FD. Stomatal development. Annu Rev Plant Biol. 2007;58:163–81. doi: 10.1146/annurev.arplant.58.032806.104023. PMID: 17201685.

46. Vercruysse J, Baekelandt A, Gonzalez N, Inzé D. Molecular networks regulating cell division during Arabidopsis leaf growth. J Exp Bot. 2020 Apr 23;71(8):2365–2378. doi: 10.1093/jxb/erz522. PMID: 31748815; PMCID: PMC7178401.

47. Kim JH. Biological roles and an evolutionary sketch of the GRF-GIF transcriptional complex in plants. BMB Rep. 2019 Apr;52(4):227–238. doi: 10.5483/BMBRep.2019.52.4.051. PMID: 30885290; PMCID: PMC6507847.

48. Li K, Yu R, Fan LM, Wei N, Chen H, Deng XW. DELLA-mediated PIF degradation contributes to coordination of light and gibberellin signalling in Arabidopsis. Nat Commun. 2016 Jun 10;7:11868. doi: 10.1038/ncomms11868. PMID: 27282989; PMCID: PMC4906400.

49. Hornitschek P, Lorrain S, Zoete V, Michielin O, Fankhauser C. Inhibition of the shade avoidance response by formation of non-DNA binding bHLH heterodimers. EMBO J. 2009 Dec 16;28(24):3893–902. doi: 10.1038/emboj.2009.306. PMID: 19851283; PMCID: PMC2797054.

50. Hao Y, Oh E, Choi G, Liang Z, Wang ZY. Interactions between HLH and bHLH factors modulate light-regulated plant development. Mol Plant. 2012 May;5(3):688–97. doi: 10.1093/mp/sss011. Epub 2012 Feb 13. PMID: 22331621; PMCID: PMC3628346.

51. Zhu L, Xin R, Bu Q, Shen H, Dang J, Huq E. A Negative Feedback Loop between PHYTOCHROME INTERACTING FACTORs and HECATE Proteins Fine-Tunes Photomorphogenesis in Arabidopsis. Plant Cell. 2016 Apr;28(4):855–74. doi: 10.1105/tpc.16.00122. Epub 2016 Apr 12. PMID: 27073231; PMCID: PMC4863390.

52. Shi H, Zhong S, Mo X, Liu N, Nezames CD, Deng XW. HFR1 sequesters PIF1 to govern the transcriptional network underlying light-initiated seed germination in Arabidopsis. Plant Cell. 2013 Oct;25(10):3770–84. doi: 10.1105/tpc.113.117424. Epub 2013 Oct 31. PMID: 24179122; PMCID: PMC3877798.

53. Xu X, Kathare PK, Pham VN, Bu Q, Nguyen A, Huq E. Reciprocal proteasome-mediated degradation of PIFs and HFR1 underlies photomorphogenic development in Arabidopsis. Development. 2017 May 15;144(10):1831–1840. doi: 10.1242/dev.146936. Epub 2017 Apr 18. PMID: 28420710; PMCID: PMC5450839.

54. Xiong F, Zhang BK, Liu HH, Wei G, Wu JH, Wu YN, Zhang Y, Li S. Transcriptional Regulation of PLETHORA1 in the Root Meristem Through an Importin and Its Two Antagonistic Cargos. Plant Cell. 2020 Dec;32(12):3812–3824. doi: 10.1105/tpc.20.00108. Epub 2020 Sep 28. PMID: 32989172; PMCID: PMC7721333.

55. Chen, D., Xu, G., Tang, W., Jing, Y., Ji, Q., Fei, Z. and Lin, R. (2013) ‘Antagonistic basic helix-loop-Helix/bZIP transcription factors form transcriptional modules that integrate light and reactive oxygen species signaling in Arabidopsis’, Plant Cell. American Society of Plant Biologists, 25(5), pp. 1657–1673. doi: 10.1105/tpc.112.104869.

56. Toledo-Ortiz G, Johansson H, Lee KP, Bou-Torrent J, Stewart K, Steel G, Rodríguez-Concepción M, Halliday KJ. The HY5-PIF regulatory module coordinates light and temperature control of photosynthetic gene transcription. PLoS Genet. 2014 Jun 12;10(6):e1004416. doi: 10.1371/journal.pgen.1004416. PMID: 24922306; PMCID: PMC4055456.

57. Dong, J., Sun, N., Yang, J., Deng, Z., Lan, J., Qin, G., He, H., Deng, X. W., Irish, V. F., Chen, H. and Wei, N. (2019) ‘The transcription factors tcp4 and pif3 antagonistically regulate organ-specific light induction of saur genes to modulate cotyledon opening during de-etiolation in arabidopsis’, Plant Cell. American Society of Plant Biologists, 31(5), pp. 1155–1170. doi: 10.1105/tpc.18.00803.

58. de Wit M, Ljung K, Fankhauser C. Contrasting growth responses in lamina and petiole during neighbor detection depend on differential auxin responsiveness rather than different auxin levels. New Phytol. 2015 Oct;208(1):198–209. doi: 10.1111/nph.13449. Epub 2015 May 11. PMID: 25963518.

59. Kidokoro S, Maruyama K, Nakashima K, Imura Y, Narusaka Y, Shinwari ZK, Osakabe Y, Fujita Y, Mizoi J, Shinozaki K, Yamaguchi-Shinozaki K. The phytochrome-interacting factor PIF7 negatively regulates DREB1 expression under circadian control in Arabidopsis. Plant Physiol. 2009 Dec;151(4):2046–57. doi: 10.1104/pp.109.147033. Epub 2009 Oct 16. PMID: 19837816; PMCID: PMC2785984.

60. Liu X, Chen CY, Wang KC, Luo M, Tai R, Yuan L, Zhao M, Yang S, Tian G, Cui Y, Hsieh HL, Wu K. PHYTOCHROME INTERACTING FACTOR3 associates with the histone deacetylase HDA15 in repression of chlorophyll biosynthesis and photosynthesis in etiolated Arabidopsis seedlings. Plant Cell. 2013 Apr;25(4):1258–73. doi: 10.1105/tpc.113.109710. Epub 2013 Apr 2. PMID: 23548744; PMCID: PMC3663266.

61. Gu D, Chen CY, Zhao M, Zhao L, Duan X, Duan J, Wu K, Liu X. Identification of HDA15-PIF1 as a key repression module directing the transcriptional network of seed germination in the dark. Nucleic Acids Res. 2017 Jul 7;45(12):7137–7150. doi: 10.1093/nar/gkx283. PMID: 28444370; PMCID: PMC5499575.

62. Weber D, Wiese C, Gessler M. Hey bHLH transcription factors. Curr Top Dev Biol. 2014;110:285–315. doi: 10.1016/B978-0-12-405943-6.00008-7. PMID: 25248480.

63. Iso T, Chung G, Hamamori Y, Kedes L. HERP1 is a cell type-specific primary target of Notch. J Biol Chem. 2002 Feb 22;277(8):6598–607. doi: 10.1074/jbc.M110495200. Epub 2001 Dec 6. PMID: 11741889.

64. Takata T, Ishikawa F. Human Sir2-related protein SIRT1 associates with the bHLH repressors HES1 and HEY2 and is involved in HES1- and HEY2-mediated transcriptional repression. Biochem Biophys Res Commun. 2003 Jan 31;301(1):250–7. doi: 10.1016/s0006-291x(02)03020-6. PMID: 12535671.

65. Gould F, Harrison SM, Hewitt EW, Whitehouse A. Kaposi’s sarcoma-associated herpesvirus RTA promotes degradation of the Hey1 repressor protein through the ubiquitin proteasome pathway. J Virol. 2009 Jul;83(13):6727–38. doi: 10.1128/JVI.00351-09. Epub 2009 Apr 15. PMID: 19369342; PMCID: PMC2698570.

66. Sang-Joo Lee, Byung Ha Lee, Jae-Hak Jung, Soon Ki Park, Jong Tae Song, Jeong Hoe Kim. GROWTH-REGULATING FACTOR and GRF-INTERACTING FACTOR Specify Meristematic Cells of Gynoecia and Anthers. Plant Physiol. 2018 Jan; 176(1): 717–729. Published online 2017 Nov 7. doi: 10.1104/pp.17.00960 PMCID: PMC5761776.

67. Reed JW, Nagpal P, Poole DS, Furuya M, Chory J. Mutations in the gene for the red/far-red light receptor phytochrome B alter cell elongation and physiological responses throughout Arabidopsis development. Plant Cell. 1993 Feb;5(2):147–57. doi: 10.1105/tpc.5.2.147. PMID: 8453299; PMCID: PMC160258.

68. Leivar P, Monte E, Oka Y, Liu T, Carle C, Castillon A, Huq E, Quail PH. Multiple phytochrome-interacting bHLH transcription factors repress premature seedling photomorphogenesis in darkness. Curr Biol. 2008 Dec 9;18(23):1815–23. doi: 10.1016/j.cub.2008.10.058. PMID: 19062289; PMCID: PMC2651225.

69. Leivar P, Monte E, Al-Sady B, Carle C, Storer A, Alonso JM, Ecker JR, Quail PH. The Arabidopsis phytochrome-interacting factor PIF7, together with PIF3 and PIF4, regulates responses to prolonged red light by modulating phyB levels. Plant Cell. 2008 Feb;20(2):337–52. doi: 10.1105/tpc.107.052142. Epub 2008 Feb 5. PMID: 18252845; PMCID: PMC2276449.

70. Yoshida Y, Sarmiento-Mañús R, Yamori W, Ponce MR, Micol JL, Tsukaya H. The Arabidopsis phyB-9 Mutant Has a Second-Site Mutation in the VENOSA4 Gene That Alters Chloroplast Size, Photosynthetic Traits, and Leaf Growth. Plant Physiol. 2018 Sep;178(1):3–6. doi: 10.1104/pp.18.00764. PMID: 30194261; PMCID: PMC6130034.

71. Gendrel AV, Lippman Z, Yordan C, Colot V, Martienssen RA. Dependence of heterochromatic histone H3 methylation patterns on the Arabidopsis gene DDM1. Science. 2002 Sep 13;297(5588):1871–3. doi: 10.1126/science.1074950. Epub 2002 Jun 20. PMID: 12077425.

72. Zhu, L. and Huq, E. (2019) ‘Characterization of Light-Regulated Protein–Protein Interactions by In Vivo Coimmunoprecipitation (Co-IP) Assays in Plants’, in Methods in Molecular Biology. Humana Press Inc., pp. 29–39. doi: 10.1007/978-1-4939-9612-4_3.

