## Supplementary Figures 1-11 for "PIF7 controls leaf cell proliferation through an AN3 substitution-repression mechanism"

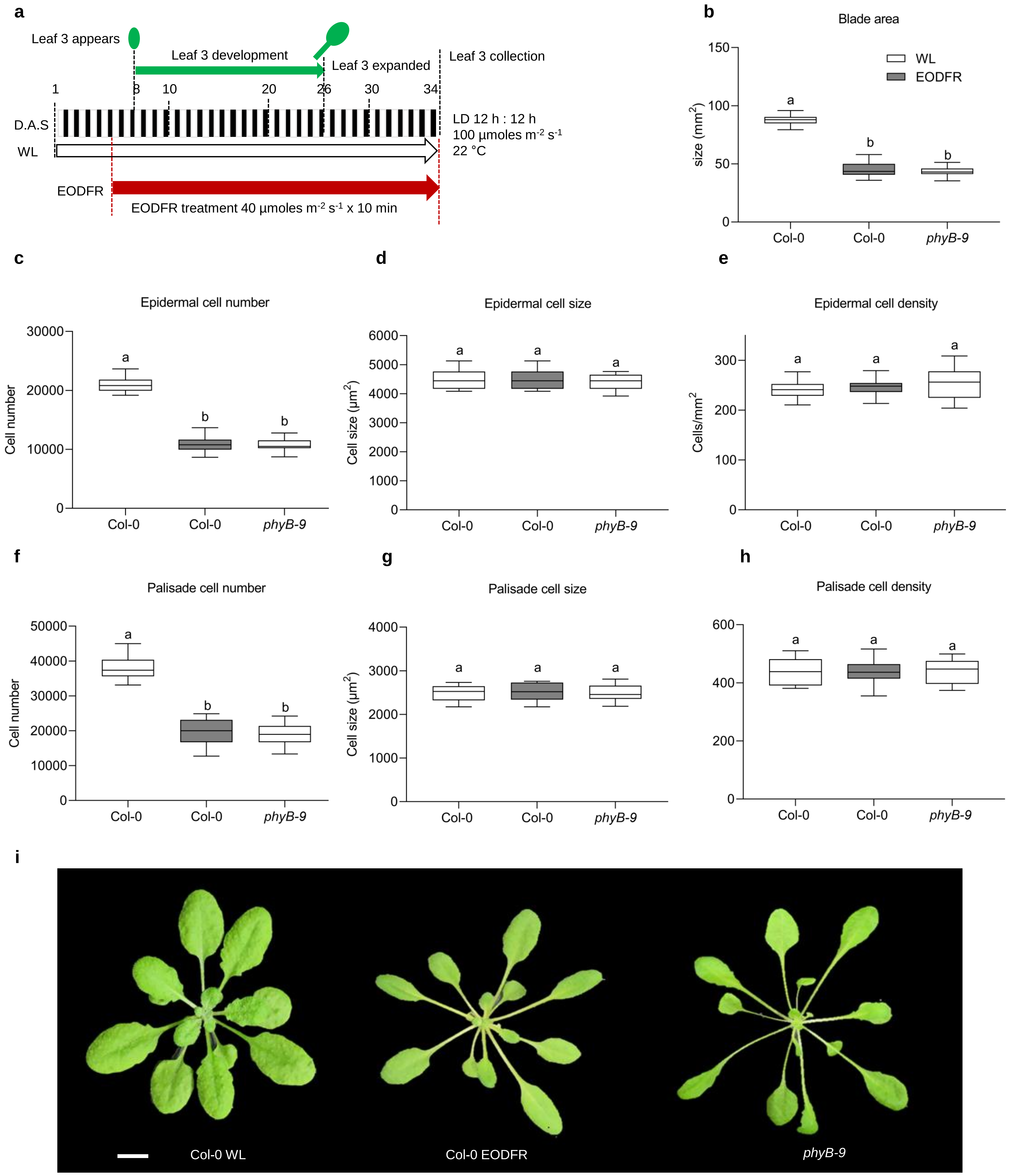


**Supplementary Fig. 1. EODFR-treated plants accurately mimics the *phyB-9* mutant phenotype. a** Schematic representation of the environmental treatment regime. White and black rectangles indicate the 12 h of day or night period, respectively. The green arrow indicates the period of L3 (leaf 3) development, which is enclosed between the two innermost black dashed lines. The plant drawn on top of day 8 indicates L3 emergence. The leaf drawn on top of day 26 indicates that L3 is fully expanded. The red dashed lines indicate the day at which a specific treatment was started and coincides with a specific coloured arrow (WL in white and EODFR in dark red). The red dashed line at the end of day 34 marks the end of each treatment. The black dashed line on day 34 indicates tissue collection**. b** Comparison of leaf blade area (ANOVA, Tukey's post hoc test, $p < 0.0001$, $n > 20$ blades). **c**, **f** epidermal and palisade cell number (ANOVA, Tukey's post hoc test, $p < 0.0001$ in (**c**, **f**), $n > 20$ blades), **d**, **g** epidermal and palisade cell size (ANOVA, Tukey's post hoc test, $p < 0.6200$ and (**d**) and $p < 0.8435$ in (**g**); 27 cells per blade, $n > 20$ blades), **e**, **h** epidermal and palisade cell density (ANOVA, Tukey's post hoc test, $p < 0.6023$ in (**e**) and $p < 0.9604$ in (h), $n > 20$ blades). **b**, **c**, **d**, **e**, **f**, **g**, **h** Error bars represent the SEM. The center of the error bars represents the mean values. Different letters denote statistically significant differences in leaf blade area, cell number, cell size and cell density between different treatments (ANOVA followed by Tukey's post hoc test). This experiment was repeated at least three times with similar results.

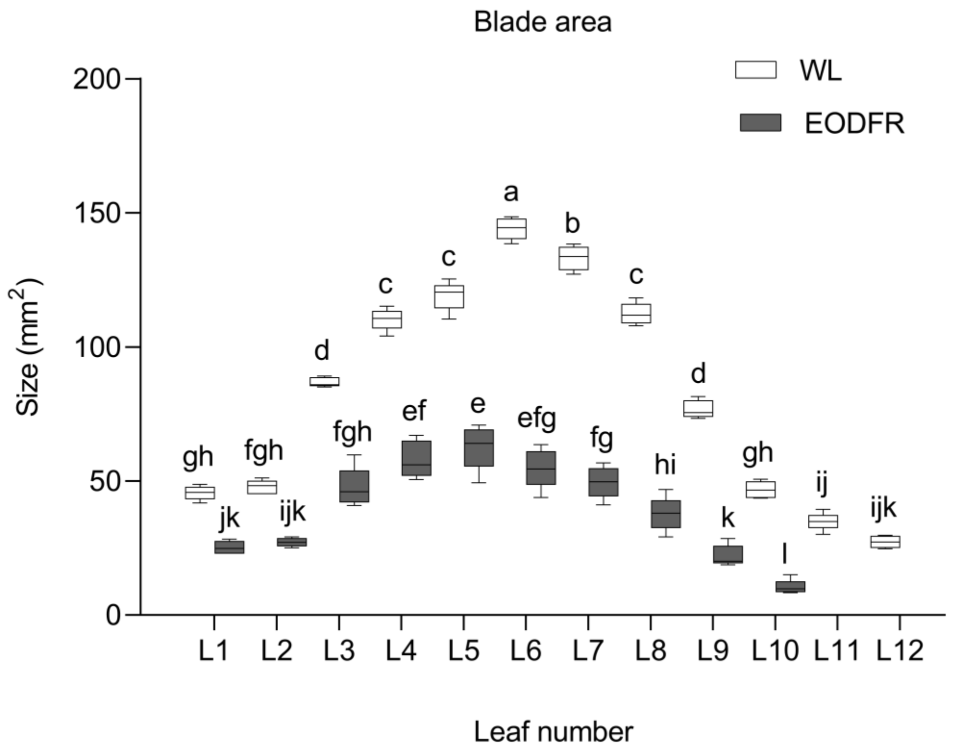


**Supplementary Fig. 2. EODFR treatment has the same effect on all rosette leaves.** Comparison of leaf blade area (ANOVA, Tukey's post hoc test, $p < 0.0001$, $n$ = 5 blades). Plants were grown under a light : dark (LD) 12 h : 12 h photoperiod, at 22 °C. On the 6th day, the plants were exposed to FR light (730 nm) at the end of each day for 10 minutes, until the 34th day. All leaf blades were harvested and measured on the 34th day. Error bars represent the SEM. The center of the error bars represents the mean values. Different letters denote statistically significant differences in leaf blade area between different treatments (ANOVA followed by Tukey's post hoc test). This experiment was repeated at least two times with similar results.

**a**

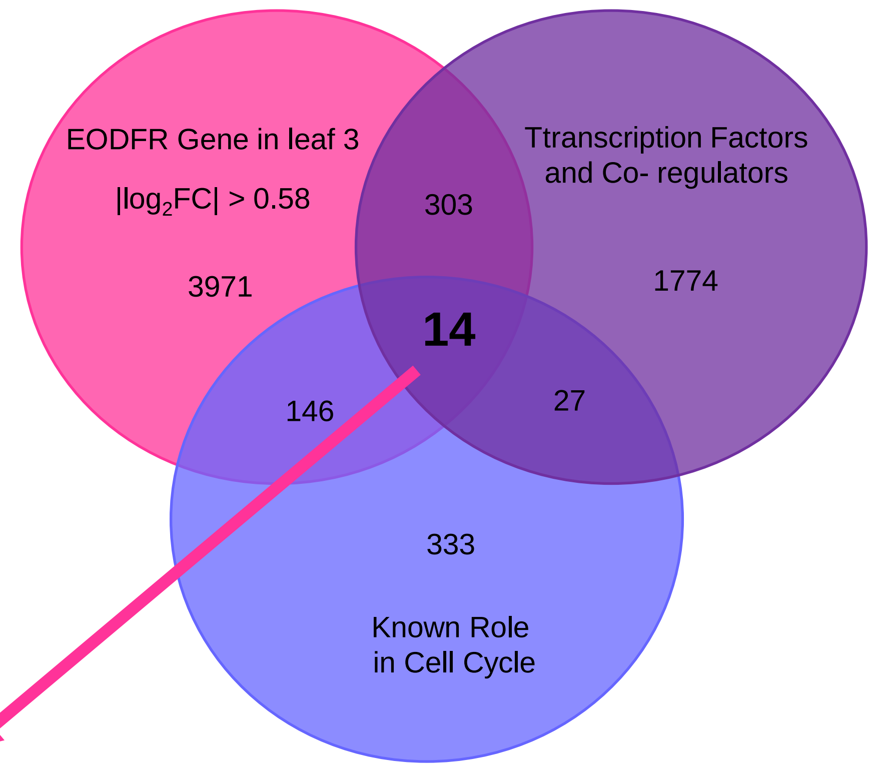


**b**

| Candidate Genes | |
|---|---|
| **GENE ID** | **Symbol** |
| AT5G28640 | *AN3* |
| AT4G37740 | *GRF2* |
| AT3G52910 | *GRF4* |
| AT2G06200 | *GRF6* |
| AT3G48160 | *DEL1* |
| AT4G37750 | *ANT* |
| AT3G01330 | *DEL3* |
| AT3G24140 | *FMA* |
| AT3G54220 | *SCR* |
| AT4G02110 | *AT4G02110* |
| AT4G32880 | *HB-8* |
| AT2G28350 | *ARF10* |
| AT2G37630 | *AS1* |
| AT5G11510 | *MYB3R-4* |

**Supplemental Fig. 3. The Venn diagram shows the overlap between candidates genes that are differentially expressed in L3 EODFR mRNAseq. a** Venn diagram of EODFR differentially expressed (left), transcription factors and co-regulators (right) and known cell cycle (bottom) genes. **b** Table showing the 14 common genes between all three data sets, representing the list of candidates. In the Venn diagrams, EODFR differentially expressed genes were extracted from our published mRNAseq (Romanowski et al., 2021), a manually curated list of transcription factors and co-regulators which were obtained from PlantTF (http://planttfdb.gao-lab.org/), AgrisTF (https://agris-knowledgebase.org/AtTFDB/) and co-activators (GO:0003713: transcription coactivator activity) and co-repressors (GO:0016564: transcription repressor activity) extracted from Virtual Plant 1.3 (Katari et al., 2010), and a list of cell cycle genes which were extracted from the Kim and kendi, 2004; Lee et al., 2009 and Van leene et al., 2010 (Supplementary Table 1).

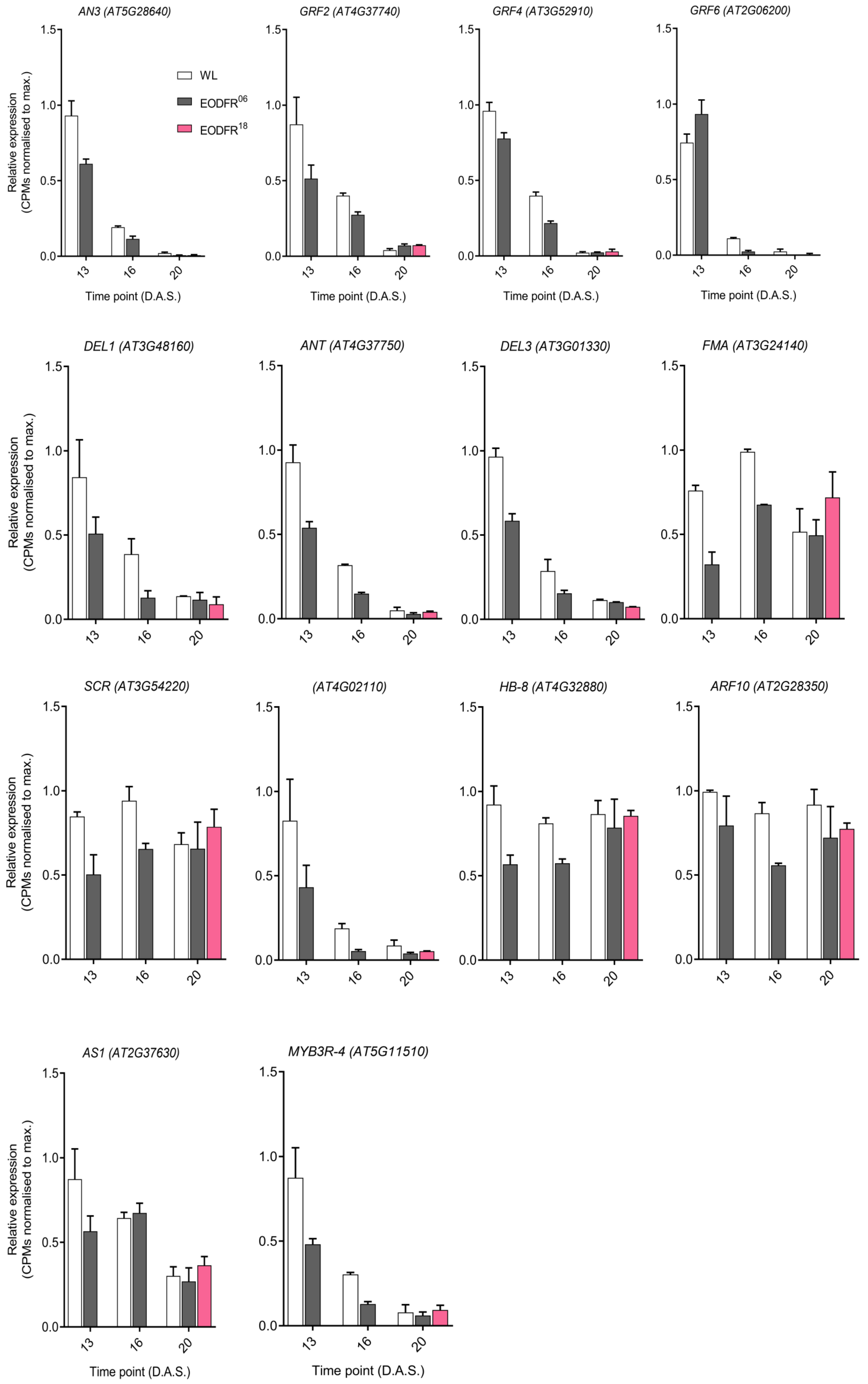


**Supplementary Fig. 3. c  Gene expression of leaf cell cycle transcriptional regulator genes from the L3 EODFR mRNAseq dataset** (Romanowski et al., 2021) shown as normalized counts relative to maximum (n = 2 biological replicates per time point per condition). Error bars represent SEM. Colour bars indicate light treatment conditions (WL (white bars), EODFR06 (grey bars) and EODFR18 (pink bars)).

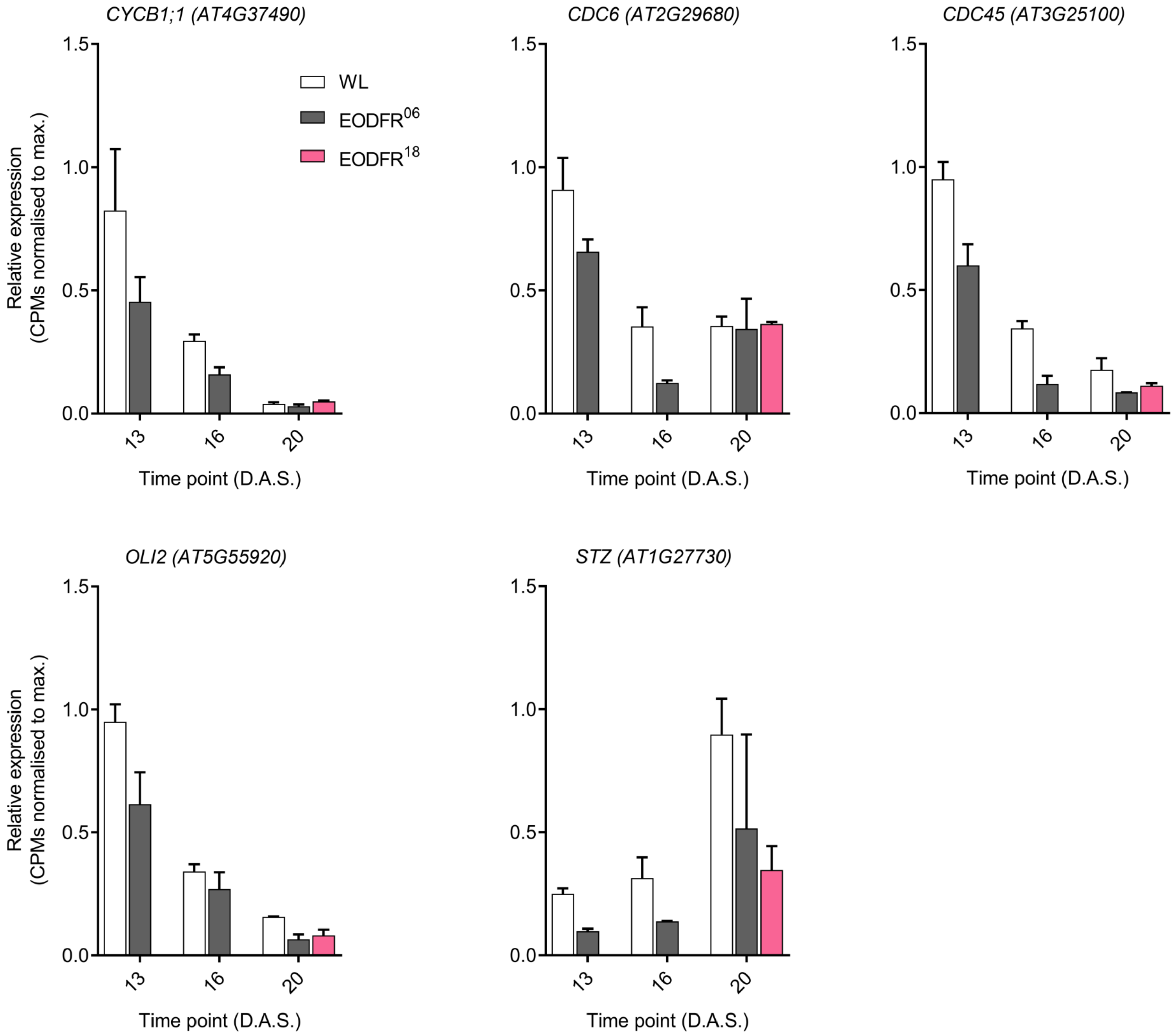


**Supplementary Fig. 3. d Gene expression of leaf cell cycle AN3 target genes from the L3 EODFR mRNAseq dataset** (Romanowski et al., 2021) shown as normalized counts relative to maximum (n = 2 biological replicates per time point per condition). Error bars represent SEM. Colour bars indicate light treatment conditions (WL (white bars), EODFR06 (grey bars) and EODFR18 (pink bars)).

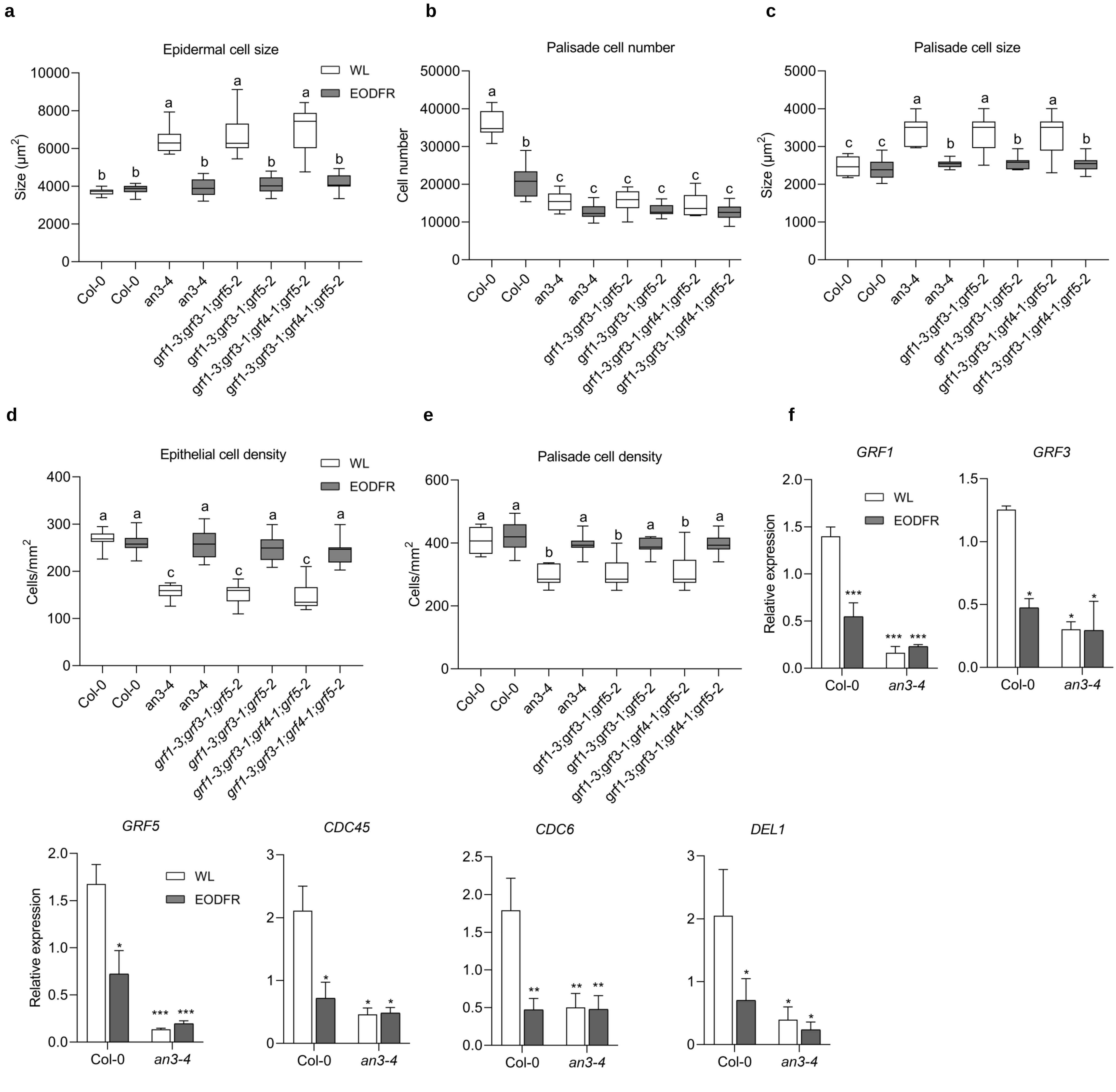


**Supplementary Fig. 4.** ***an3*** **and** ***grf*** **mutants shows no change in cell number in EODFR condition and EODFR regulated AN3 target genes transcript levels are constitutively low in** ***an3*** **blades.** **a** Epidermal cell size (ANOVA, Tukey's post hoc test, $p < 0.0001$; 27 cells per blade, $n = 15$ blades). **b** palisade cell number (ANOVA, Tukey's post hoc test, $p < 0.0001$, $n = 15$ blades), **c** palisade cell size (ANOVA, Tukey's post hoc test, $p < 0.0001$; 27 cells per blade, $n = 15$ blades) and **d, e** epidermal and palisade cell density (ANOVA, Tukey's post hoc test, $p < 0.0001$ in (**d**, **e**); $n = 15$ blades). **a**, **b, c, d, e** Error bars represent the SEM. The center of the error bars represents the mean values. Different letters denote statistically significant differences in leaf blade area, cell number, cell size and cell density between genotypes and different treatments (ANOVA followed by Tukey's post hoc test). This experiment was repeated at least two times with similar results. **f** *GRF1, GRF3, GRF5, CDC45, CDC6* and *DEL1* messenger RNA (mRNA) level using quantitative reverse transcription (RT-qPCR) after the shift from white light to EODFR in Col-0 and *an3-4*. Seedlings were grown for 13 days under a light : dark (LD) 12 h : 12 h photoperiod, at 22 °C. On day 13, seedlings were either shifted to EODFR treatment or kept in the white light control condition. L3 blades were harvested 13 days after sowing at zeitgeber (ZT) 24. The transcript levels were calculated relative to those of *PP2A*. Error bars represent the s.d. of three biological replicates. The center of the error bars represents the mean values. (*$p < 0.05$, **$p < 0.01$ and ***$p < 0.001$ , Student's *t*-test).

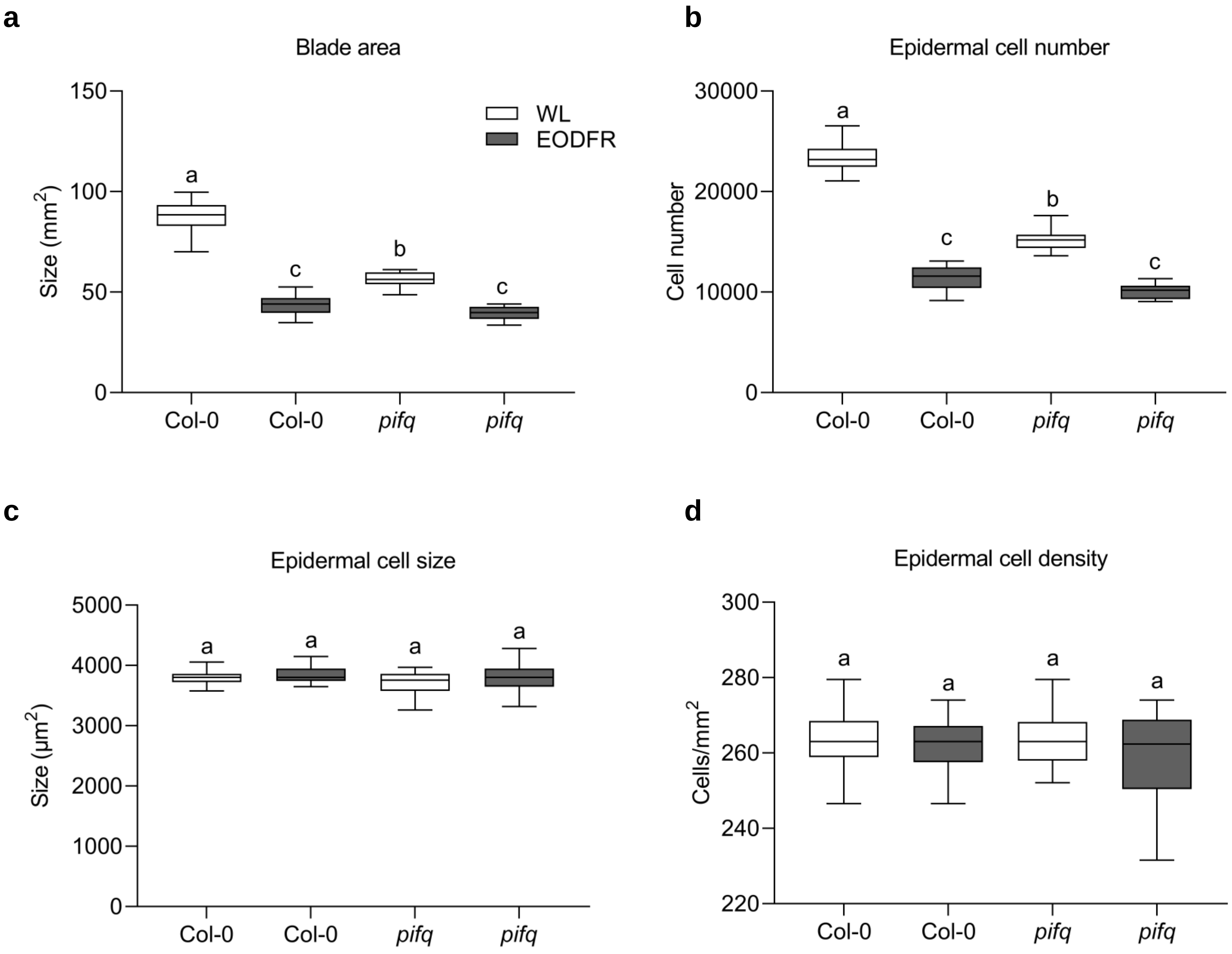


**Supplementary Fig. 5. The *pifq* mutant still reduces leaf cell number in response to EODFR treatment. a** Comparison of leaf blade area (ANOVA, Tukey's post hoc test, $p < 0.0001$, $n > 20$ blades). **b** epidermal cell number (ANOVA, Tukey's post hoc test, $p < 0.0001$, $n > 20$ blades), **c** epidermal cell size (ANOVA, Tukey's post hoc test, $p = 0.1175$; 27 cells per blade, $n > 20$ blades), **d** epidermal cell density (ANOVA, Tukey's post hoc test, $p = 0.2315$, $n > 20$ blades). Plants were grown under a light : dark (LD) 12 h : 12 h photoperiod, at 22 °C. On the 6th day, the plants were exposed to EODFR treatment, until the 34th day. All leaf 3 blades were harvested and measured on the 34th day. **a**, **b**, **c**, **d** Error bars represent the SEM. The center of the error bars represents the mean values. Different letters denote statistically significant differences in leaf blade area, cell number, cell size and cell density between different treatments (ANOVA followed by Tukey's post hoc test). This experiment was repeated at least two times with similar results.

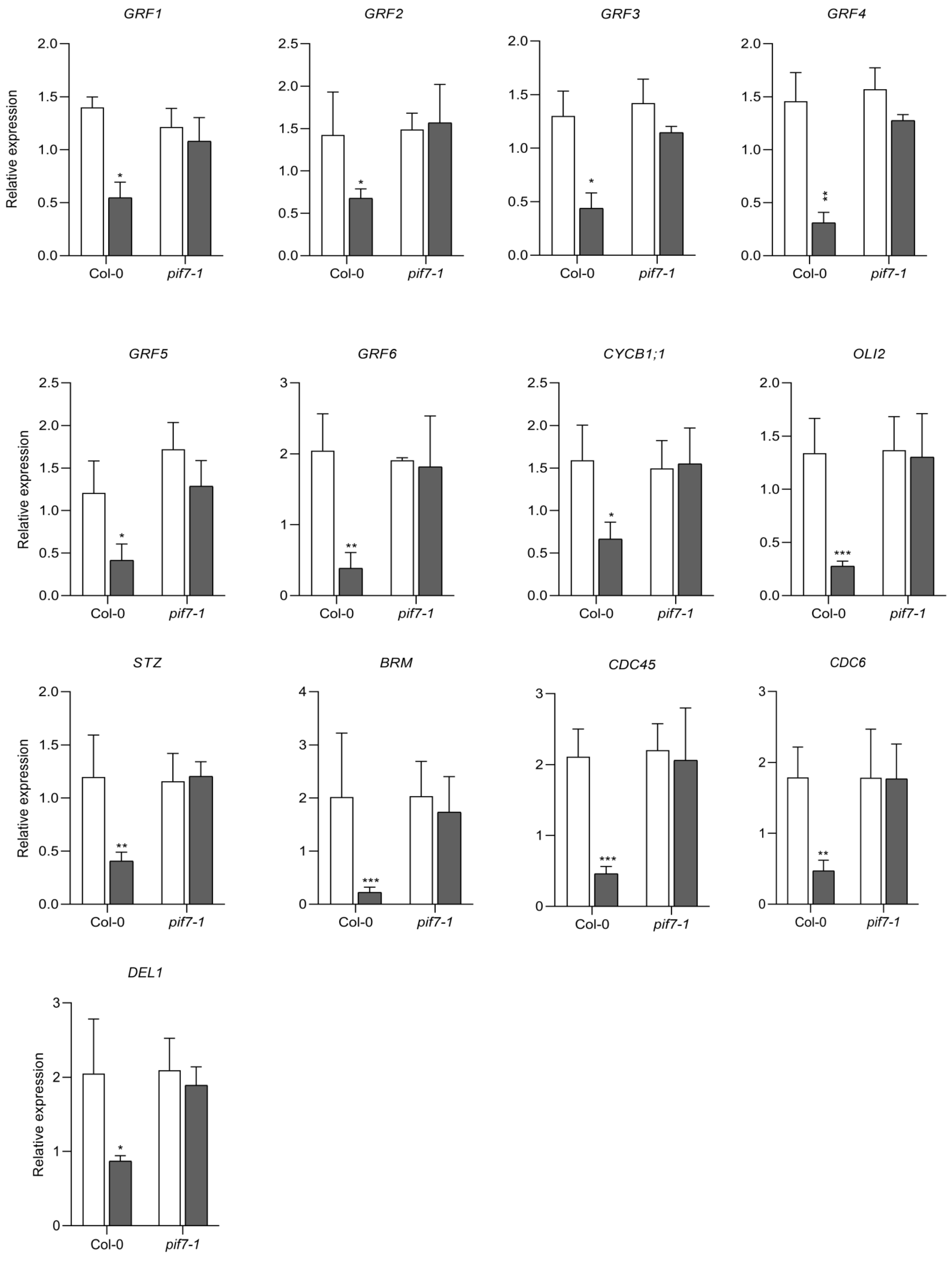


**Supplementary Fig. 6. PIF7 regulates the transcription level of AN3 target genes in EODFR.** *GRF1, GRF2, GRF3, GRF4, GRF5, GRF6, CYCB1;1, OLI2, STZ, BRM, CDC45, CDC6* and *DEL1* messenger RNA (mRNA) level using quantitative reverse transcription (RT-qPCR) after the shift from white light to EODFR in Col-0 and *pif7-1*. Seedlings were grown for 13 days under a light : dark (LD) 12 h : 12 h photoperiod, at 22 °C. On day 13, seedlings were either shifted to EODFR treatment or kept in control white light condition. L3 blades were harvested 13 days after sowing at zeitgeber (ZT) 24. The transcript levels were calculated relative to those of *PP2A*. Error bars represents the s.d. of three biological replicates. The center of the error bars represents the mean values. (*$p < 0.05$, **$p < 0.01$ and ***$p < 0.001$, Student's *t*-test).

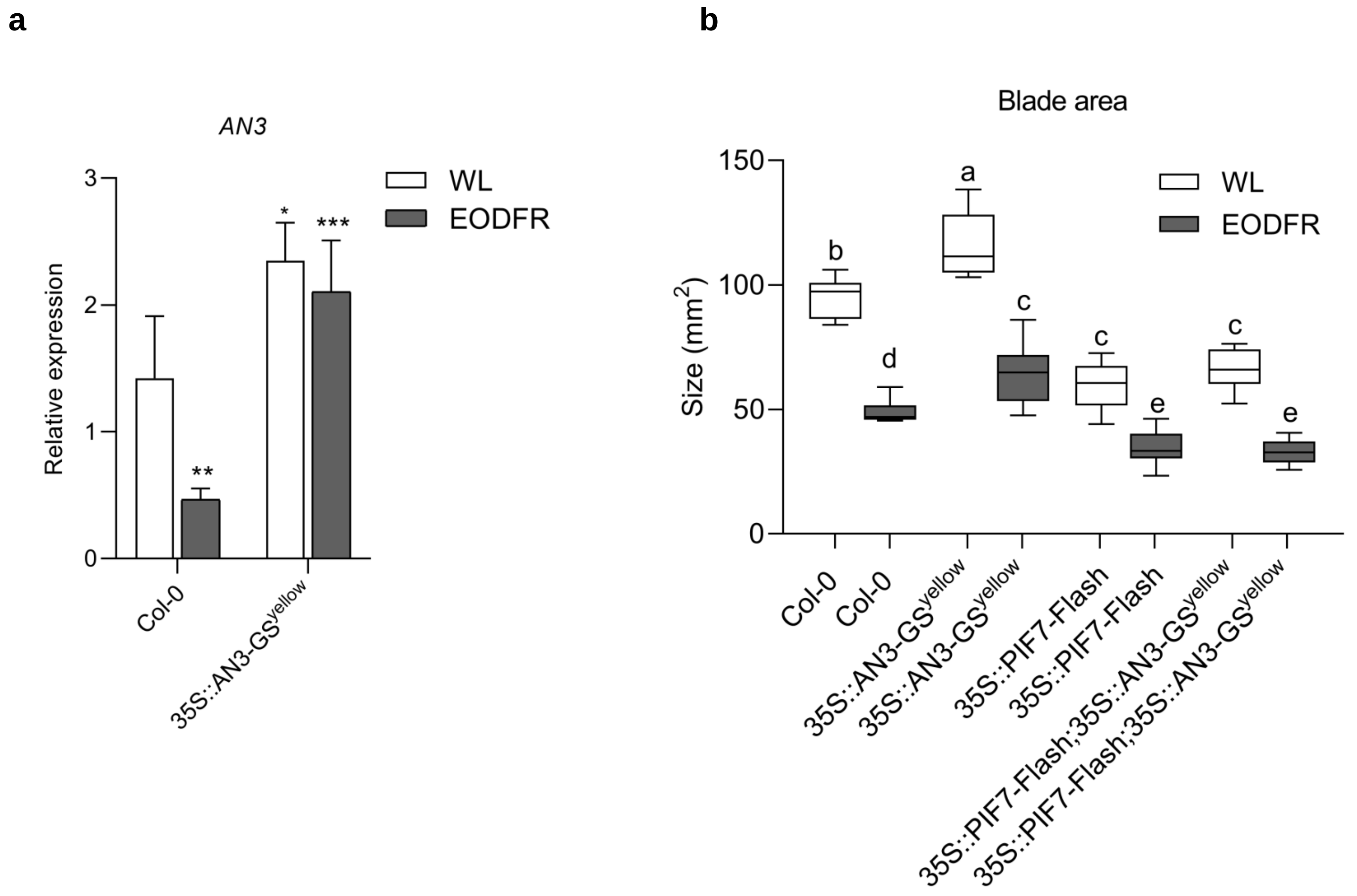


**Supplementary Fig. 7. Overexpression of *PIF7* shows dominant effect on AN3 during leaf expansion. a** *AN3* messenger RNA (mRNA) level using quantitative reverse transcription (RT-qPCR) after the shift from white light to EODFR in Col-0 and *35S::AN3-GS*$^{yellow}$. Seedlings were grown for 13 days under a light : dark (LD) 12 h : 12 h photoperiod, at 22 °C. On day 13, seedlings were either shifted to EODFR treatment or kept in control white light condition. L3 blades were harvested 13 days after sowing at zeitgeber (ZT) 24. The transcript levels were calculated relative to those of *PP2A*. Error bars represents the s.d. of three biological replicates. The center of the error bars represents the mean values. (*$p < 0.05$, **$p < 0.01$ and ***$p < 0.001$, Student's *t*-test). **b** Comparison of leaf blade area (ANOVA, Tukey's post hoc test, $p < 0.0001$, $n > 12$ blades), between Col-0 and 35S::AN3-GS$^{yellow}$, 35S::PIF7-Flash and 35S::AN3-GS$^{yellow}$; 35S::PIF7-Flash plants. Plants were grown under a light : dark (LD) 12 h : 12 h photoperiod, at 22 °C. On the 6th day, the plants were exposed to EODFR treatment, until the 34th day. All leaf 3 blades were harvested and measured on the 34th day. For (**b),** Error bars represents the SEM. The center of the error bars represents the mean values. Different letters denote statistically significant differences in leaf blade area between different treatments (ANOVA followed by Tukey's post hoc test). This experiment was repeated at least three times with similar results.

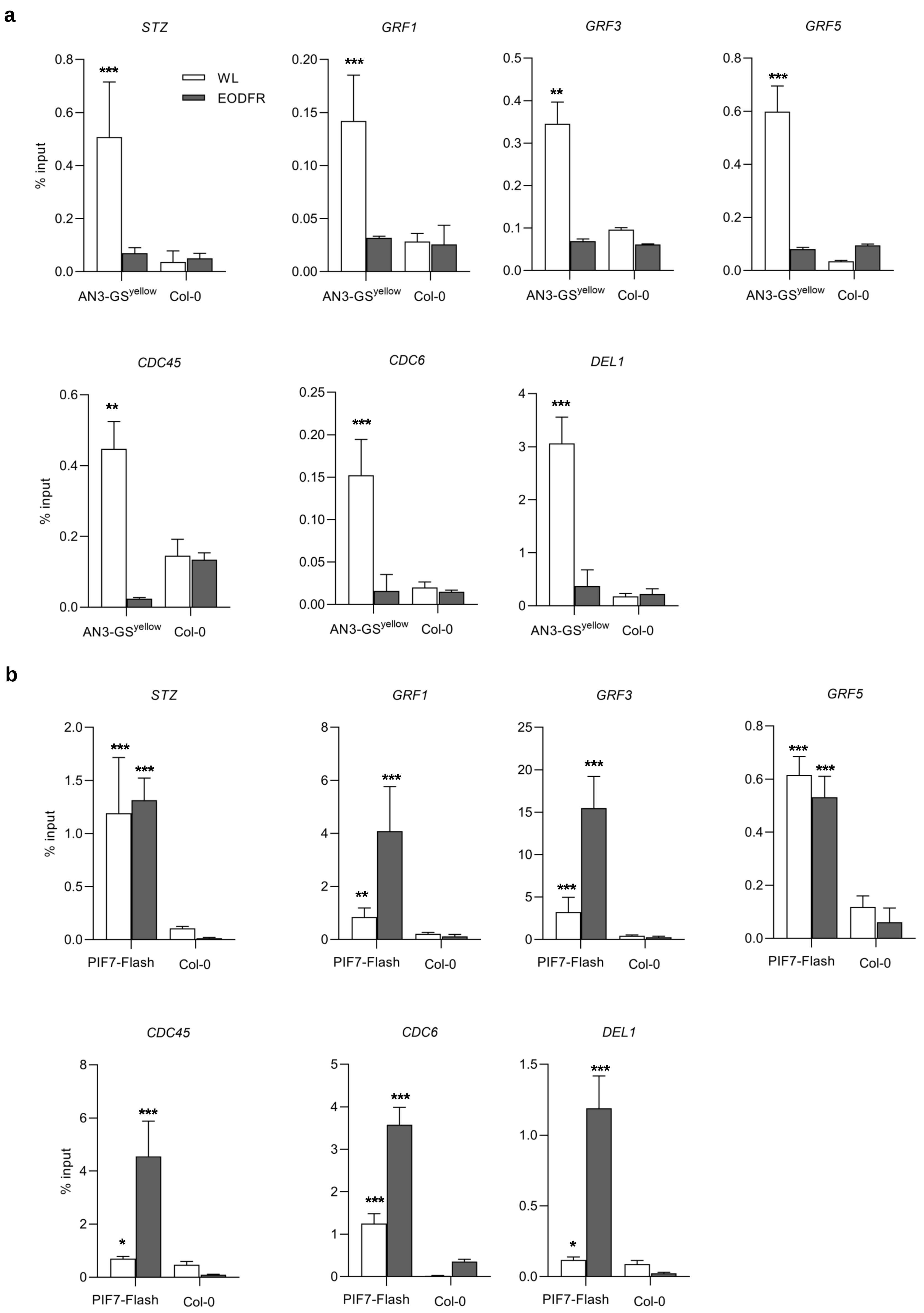


**Supplementary Fig. 8. PIF7 replaces AN3 and directly binds to the target gene promoters of cell cycle genes.**

**a, b** ChIP assay showing enrichment of 35S::AN3-Gs$^{yellow}$ and 35S::PIF7-Flash on *STZ, GRF1, GRF3, GRF5, CDC45, CDC6* and *DEL1* promoter fragments containing the G-box or PBE motif. Seedlings were grown under a light : dark (LD) 12 h : 12 h photoperiod, at 22 °C. On day 13, seedlings were treated with EODFR or kept in the white light control condition. Samples were taken at ZT14 two hours after the EODFR treatment. The 35S::AN3-Gs$^{yellow}$ and 35S::PIF7-Flash were incubated with anti-GFP and anti-Myc antibodies, respectively, and precipitated by Dynabeads protein A. The enrichment of fragments was determined by qPCR. The wild type (Col-0) plants acted as a control. Error bars represent the s.d. of three biological replicates. The center of the error bars represent the mean values. (*$p < 0.05$, **$p < 0.01$ and ***$p < 0.001$, Student's *t*-test).

**c**

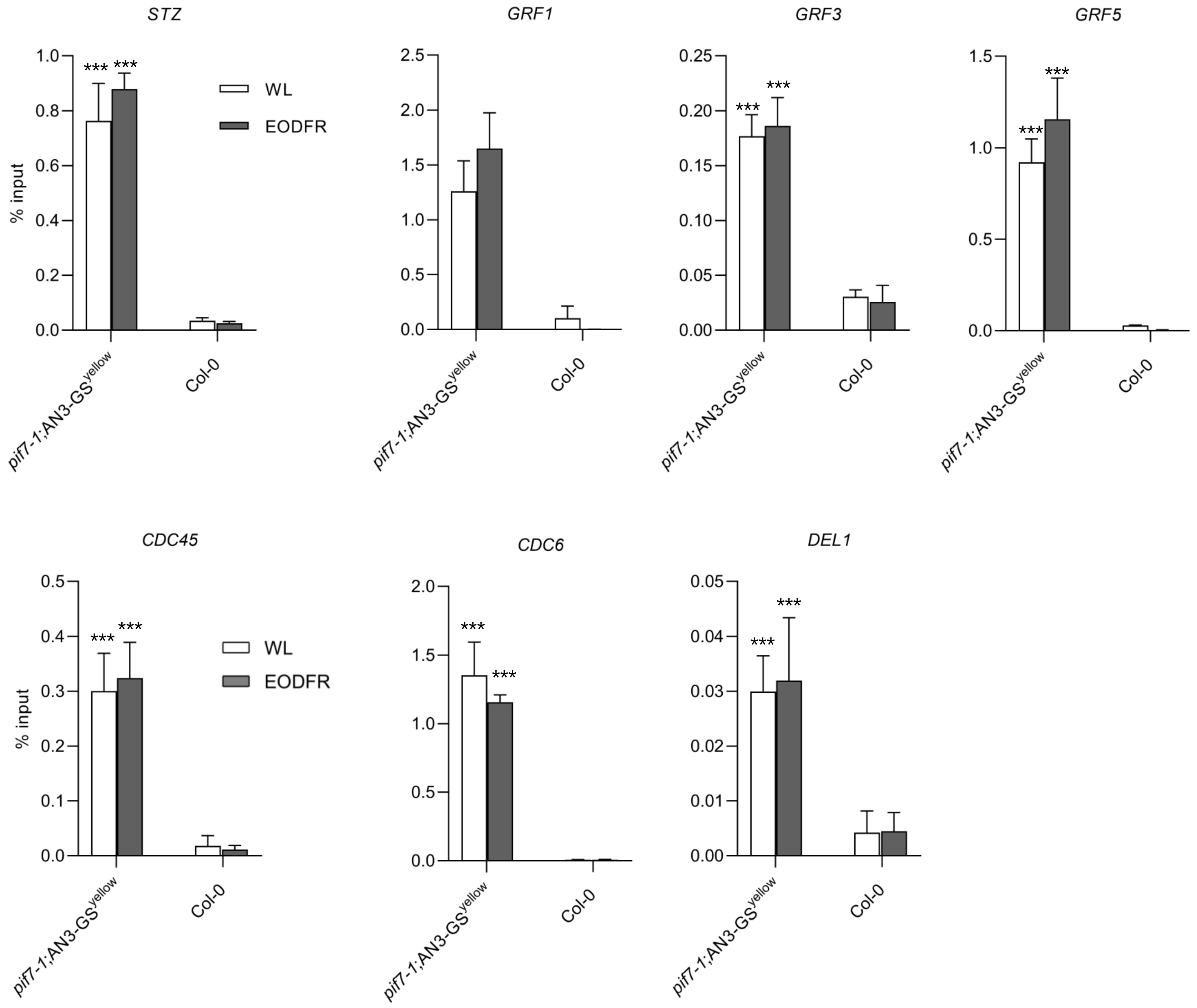


**Supplementary Fig. 8. PIF7 replaces AN3 and directly binds to the target gene promoters of cell cycle genes.**

**c** ChIP assay showing enrichment of *pif7-1*;35S::AN3-GS$^{yellow}$ on *STZ, GRF1, GRF3, GRF5, CDC45, CDC6* and *DEL1* promoter fragments containing the G-box or PBE motif. Seedlings were grown under a light : dark (LD) 12 h : 12 h photoperiod, at 22 °C. On day 13, seedlings were treated with EODFR or kept in the white light control condition. Samples were taken at ZT14 two hours after the EODFR treatment. The *pif7-1;*35S::AN3-GS$^{yellow}$ were incubated with anti-GFP and precipitated by Dynabeads protein A. The enrichment of fragments was determined by qPCR. The wild type (Col-0) plants acted as a control. Error bars represent the s.d. of three biological replicates. The center of the error bars represent the mean values. (*$p < 0.05$, **$p < 0.01$ and ***$p < 0.001$, Student's *t*-test).

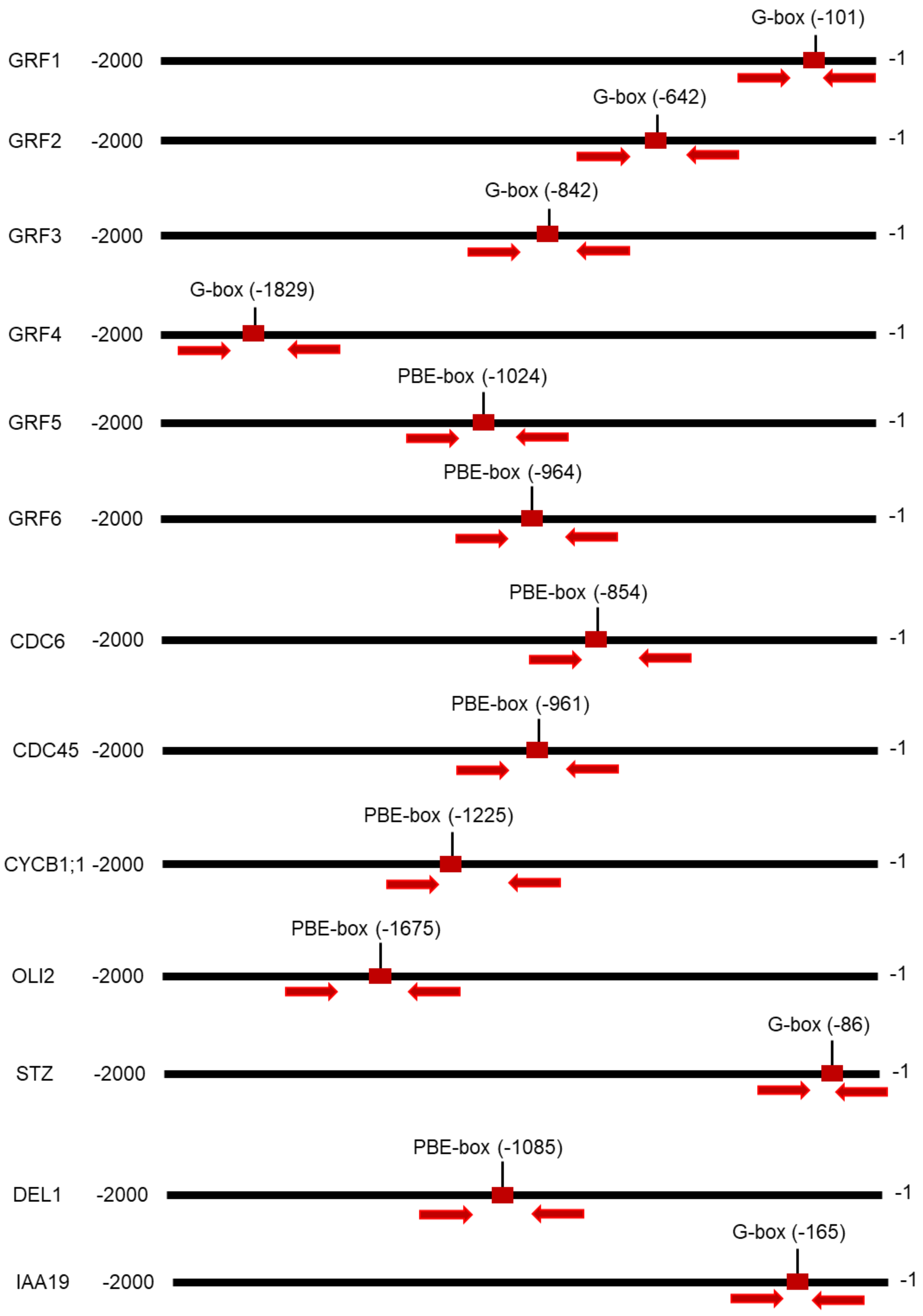


**Supplementary Fig. 9. Schematic representation of putative AN3 target genes promoters which also harbour putative PIF7 binding elements (G-box or PBE box).** A 2-kb upstream promoter region of each target gene obtained with PlantPAN 3.0 (http://plantpan.itps.ncku.edu.tw/) is shown for each gene. G-box (CACGTG) and PBE-box (CA[TG/CA]TG target binding sites appear as red boxes. The arrow pairs indicate the fragments amplified by ChIP-qPCR in this study. Numbers represent the position in base pairs upstream from the transcription start site. The numbers in between parenthesis indicate the position of the G-box or PBE-box upstream from the transcription start site.

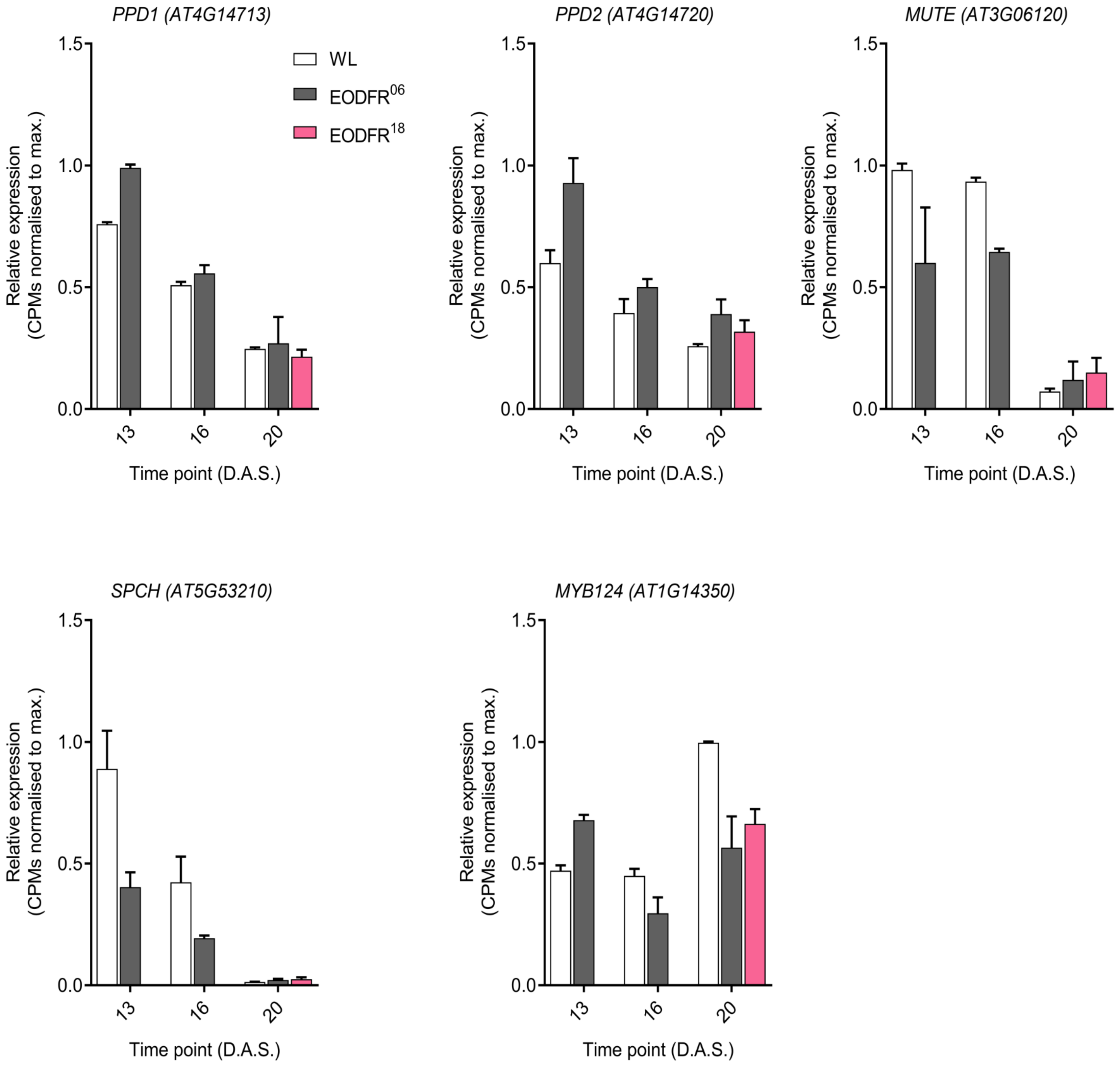


**Supplementary Fig. 10. Gene expression of meristemoid phase of cell proliferation genes from the L3 EODFR mRNAseq.** The Transcript levels are shown as normalized counts relative to maximum (n = 2 biological replicates per time point per condition) (Romanowski et al., 2021). Colours of bars indicates (WL (white bars), EODFR06 (grey bars) and EODFR18 (pink bars)) light treatment conditions. Error bars represent S.E.M. L3 = Leaf 3; D.A.S. = Days After Sowing; WL = white light; EOD = end of day; FR = far-red; $EODFR^{06}$ = EODFR since day 6; $EODFR^{18}$ = EODFR since day 18.

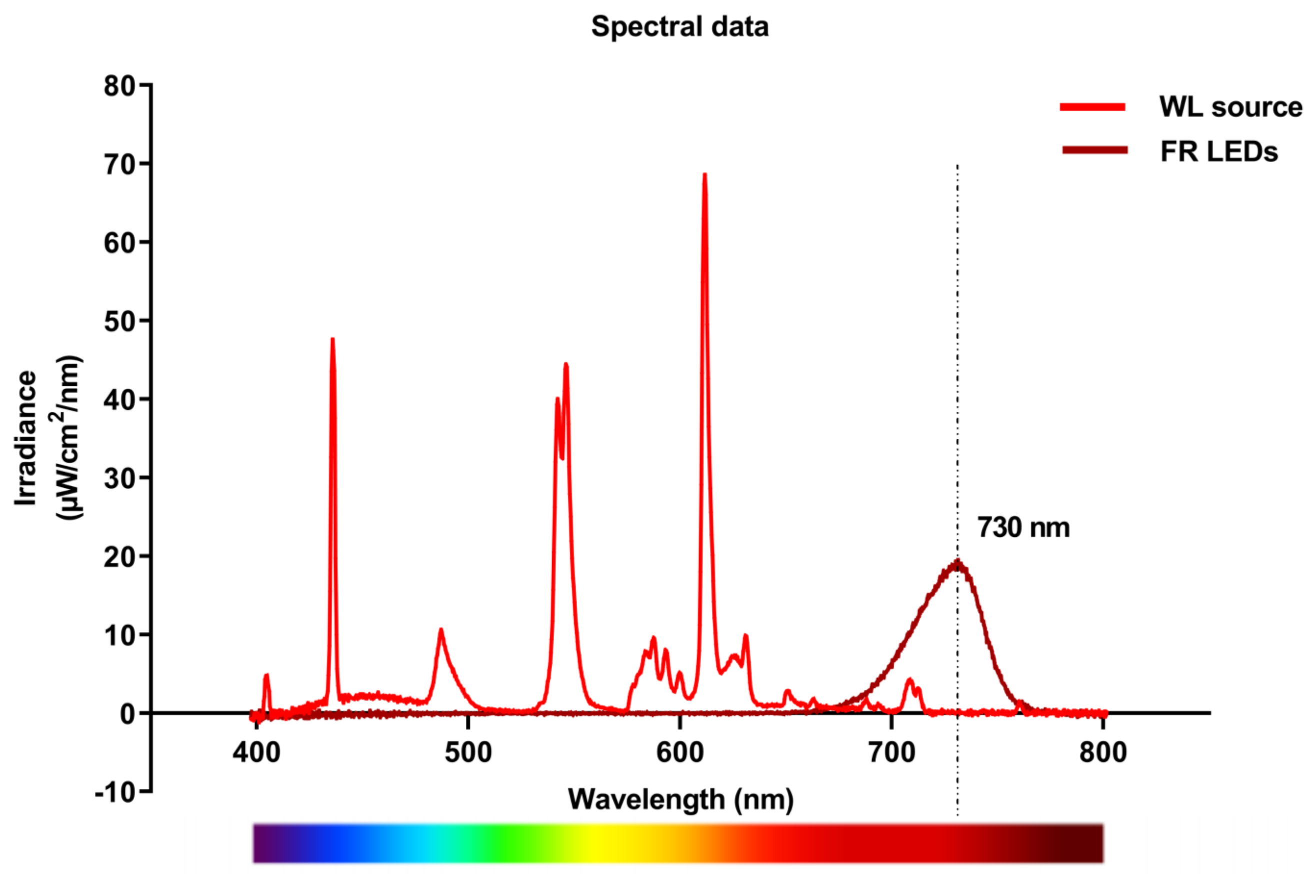


**Supplemental Fig. 11. Spectral data information used in this study.** Irradiance plot for the white light (WL) source (red line) and far-red (FR) LEDs (dark red line) used in this work. The dashed line marks the peak of FR emission (730 nm).

**References**

1. Katari MS, Nowicki SD, Aceituno FF, Nero D, Kelfer J, Thompson LP, Cabello JM, Davidson RS, Goldberg AP, Shasha DE, Coruzzi GM, Gutiérrez RA. VirtualPlant: a software platform to support systems biology research. Plant Physiol. 2010 Feb;152(2):500-15. doi: 10.1104/pp.109.147025. Epub 2009 Dec 9. PMID: 20007449; PMCID: PMC2815851.
